# PTPN22 Regulates T Cell Synapse Formation through PSTPIP1- Dependent Actin Remodeling

**DOI:** 10.1101/2025.04.24.650379

**Authors:** Megan D. Joseph, Cecilia Zaza, Olivia P. L. Dalby, Michael L. Dustin, Andrew P. Cope, Sabrina Simoncelli

## Abstract

Protein tyrosine phosphatase non-receptor type 22 (PTPN22) is a critical regulator of T cell signaling, working in concert with C-terminal SRC kinase (Csk) to dephosphorylate key signaling proteins and suppress lymphocyte activation. The R620W variant of PTPN22, one of the most prevalent mutations associated with autoimmune diseases, has been implicated in altered T cell responses, although its broader effects on T cell activation are not fully understood. Recent studies have uncovered a novel interaction between PTPN22 and proline-serine-threonine phosphatase interacting protein 1 (PSTPIP-1), a cytoskeletal adaptor protein involved in F-actin remodeling. PSTPIP-1 recruits Wiskott-Aldrich Syndrome Protein (WASp) to facilitate actin foci formation, a process integral to stabilizing TCR microclusters and amplifying downstream signaling. Given that mutations in PSTPIP-1 impair actin remodeling and localize within the PTPN22 binding domain, we hypothesized that PTPN22 modulates actin dynamics through its interaction with PSTPIP-1. Using live and fixed multi-color super-resolution fluorescence imaging, we demonstrate that PTPN22 deficiency or inhibition of its phosphatase activity leads to aberrant Arp2/3-dependent actin remodeling and exaggerated calcium signaling, particularly under low-affinity TCR stimulation. Single-protein resolution imaging via DNA-PAINT further revealed disrupted nanoscale clustering of PSTPIP-1 and TCR in PTPN22-deficient T cells, uncovering a previously unrecognized PSTPIP-1–TCR interaction in the absence of PTPN22. These findings highlight a novel PTPN22–PSTPIP-1 signaling axis, offering new insights into the molecular mechanisms that may contribute to autoimmune disease susceptibility.

**One Sentence Summary:** PTPN22 and PSTPIP-1 control actin remodeling upon T cell synapse formation, modulating TCR clustering and activation.

## INTRODUCTION

Protein tyrosine phosphatase non-receptor type 22 (PTPN22), also known as lymphoid tyrosine phosphatase (Lyp) in humans, plays a central role in modulating immune signaling through its phosphatase activity (*1*). A genetic variant in the PTPN22 coding region (PTPN22-R620W), is strongly linked to a wide range of autoimmune diseases including systemic lupus erythematosus (SLE), rheumatoid arthritis (RA), and type 1 diabetes (T1D)(*1–5*). This mutation disrupts T and B cell activation and signaling (*6*). PTPN22’s established role in T cell activation centers on its interaction with Csk, a negative regulator of TCR signaling. Csk and PTPN22 operate in tandem to deactivate src and syk kinase substrates upon T cell activation. Specifically, PTPN22 dephosphorylates the activatory Tyr394 on Lck, while Csk phosphorylates the inhibitory Tyr505 of Lck(*7*, *8*). In fact, super-resolution microscopy studies have highlighted PTPN22’s crucial role in dynamically re-recruiting Csk to membrane-proximal regions during mature T cell synapses, a process essential for Csk’s downregulatory function (*9*).

Recent interactome analysis has reshaped our understanding of PTPN22 function, identifying Proline-Serine-Threonine Phosphatase Interacting Protein 1 (PSTPIP-1) as its primary constitutive binding partner—a key cytoskeletal-associated protein that orchestrates actin remodeling during T cell activation (*10–12*). This discovery expands PTPN22’s known role beyond TCR signaling, positioning it as a potential regulator of cytoskeletal dynamics and immune synapse organization. PSTPIP-1 associates with the actin-regulatory protein Wiskott-Aldrich syndrome protein (WASp), which facilitates the nucleation of branched actin networks through the Arp2/3 complex (*13*, *14*). Although less studied than other actin modulators, PSTPIP-1’s interactions with CD2 and actin-binding proteins suggest its role in positioning actin nucleators, including WASp, at sites of TCR engagement (*13*, *15*, *16*). These interactions are essential for recruiting the Arp2/3 complex, promoting actin polymerization, and forming branched actin networks at the T cell immune synapse, critical processes for stabilizing TCR clusters and enhancing TCR signaling (*16*). Further supporting the role of PTPN22 in cytoskeletal regulation, its interaction with PSTPIP-1 is mediated through the carboxyl-terminal homology (CTH) domain of PTPN22 and the F-bar domain of PSTPIP-1 (*11*). Notably, mutations within PSTPIP-1’s F-bar domain have been linked to PSTPIP- 1-associated inflammatory diseases (PAIDs), a group of autoinflammatory disorders that disrupt actin formation in T cells (*17–19*).

The discovery of the PTPN22–PSTPIP-1 interaction unveils a previously unrecognized mechanism linking phosphatase signaling to cytoskeletal remodeling during T cell activation. Investigating this interaction could yield critical insights into immune regulation and reveal how its dysregulation contributes to diseases associated with PTPN22 and PSTPIP-1 mutations. To dissect the functional significance of the PTPN22–PSTPIP-1 interaction, we employed live and fixed super-resolution fluorescence microscopy to examine actin remodeling in wild-type (WT) and PTPN22 knockout (KO) Jurkat T cells upon activation. Our findings establish PTPN22 as a key regulator of actin dynamics by controlling PSTPIP-1 localization and function, which directly impacts TCR clustering and immune synapse formation. Notably, PTPN22-dependent actin remodeling requires phosphatase activity and is mediated by the Arp2/3 complex. Collectively, our study uncovers a novel mechanism in which PTPN22 modulates actin remodeling and TCR organization through PSTPIP-1, providing crucial insights into immune regulation and potential therapeutic targets for autoimmune diseases and cancer immunotherapy.

## RESULTS

### PTPN22 Deficiency Alters Actin Remodeling During Immunological Synapse Formation

We investigated the role of PTPN22 in actin remodeling during T cell activation using an isogenic human T cell line lacking PTPN22, generated via CRISPR gene editing and validated in previous work (*20*). Visualization of cytoskeletal rearrangements during immunological synapse formation was achieved by labeling live WT and PTPN22 KO Jurkat T cells with SiR-actin and exposing them to aCD3/aCD28-coated microscope coverslips. Actin dynamics at the cell contact interface were captured using super-resolution fluorescence microscopy with optical pixel reassignment (SoRa) at 10-second intervals, from initial contact up to 20 minutes (see Movie S1 and S2). This approach enabled fast, high-resolution imaging of F-actin (203 ± 45 nm laterally, as calculated by Fourier Ring Correlation analysis, (*21*)) without high photodamage or bleaching in live samples.

Figure 1A presents representative images of F-actin remodeling in WT and PTPN22 KO Jurkats over time following aCD3/aCD28 activation. This stimulation induced cell adhesion, actin remodeling, and dynamic spreading, culminating in distinct actin structures by 10 minutes. WT Jurkat T cells exhibited three distinct F-actin zones upon aCD3/aCD28 stimulation: (i) an outer dense ring corresponding to the dSMAC, (ii) a middle ring with concentric actin arcs in the pSMAC, and (iii) a central actin-depleted cSMAC (Fig. 1A, t = 20 min). In contrast, PTPN22 KO cells displayed altered actin structures, with increased central F-actin concentration around the cSMAC and a thinner peripheral branched actin ring in the dSMAC. Despite differences in actin architecture, both WT and PTPN22 KO Jurkat T cells formed similar-sized immune synapse contacts upon TCR stimulation, reaching maximum contact area around 250 seconds post-stimulation (Fig. 1B, solid lines). In contrast, neither WT nor KO cells spread in the absence of activating ligands (Fig. S1, Fig. 1B, dashed lines). The final contact area (Fig. 1C) showed no significant differences between WT and KO cells, although both displayed larger contact areas compared to unstimulated conditions. Similarly, kymograph analysis (Fig. 1D) revealed clear inward motion of F-actin under activation, flowing at comparable rates for WT ((3 ± 1) x 10^−2^ μm/min) and PTPN22 KO ((5 ± 5) x 10^−2^ μm/min) cells (Fig. 1E). Centripetal actin flow was not observed under unstimulated conditions.

**Fig. 1.**
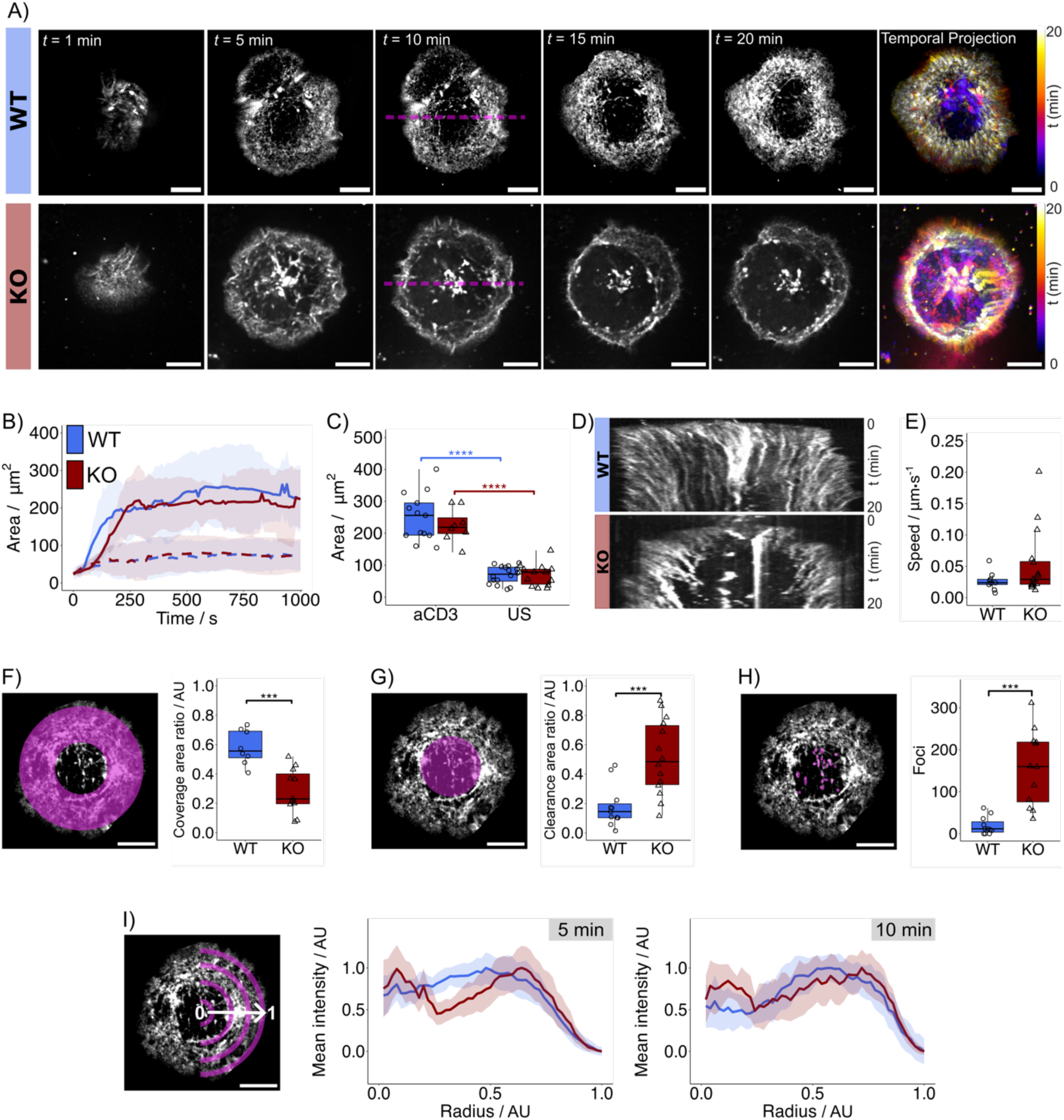
Actin remodeling in WT and PTPN22 KO Jurkat T cells interacting with aCD3/aCD28 coated slides. **(A)** Spinning disk confocal SoRa fluorescent imaging time-lapse of Jurkat T cells incubated with SiR actin from 1-20 minutes post settling. 1-, 5-, 10-, 15-, and 20- minute time frames are shown. Temporal projections of spreading are depicted with pseudo-color images with color gradients transitioning from purple to white representing early to late time- points. Scale bar = 5 μm. **(B)** Area of cell spreading over time on glass slides coated with aCD3/aCD28 (solid lines) compared to unstimulated cells (dashed lines). **(C)** Maximum area of cell spreading from **B**. **(D)** Kymographs of aCD3/aCD28 activated WT and PTPN22 KO Jurkat T cells drawn from line selection in **A**. **(E)** Initial speed of F-actin flow calculated from kymographs in **D**. **(F)** Ratio of F-actin ring coverage area depicted in purple to total cell area. **(G)** Ratio of central F-actin clearance area depicted in purple to total cell area. **(H)** The number of F-actin foci like structures found in the central regions of cells, as highlighted in purple. **(I)** Radial analysis depicting mean relative F-actin intensity from the central region (radius = 0) to the edge of the cell (radius = 1) as shown in purple. Radial analyses of cells settling on aCD3/aCD28 coated glass slides after 5 minutes and after 10 minutes are shown with grey boxes indicating time point. Solid line indicates the mean with shaded areas representing standard deviation. For all plots, n = 10 cells from 3 independent passages. Statistical tests consisted of parametric ANOVA with pairwise Tukey’s HSD for normally distributed data and non-parametric Wilcoxon signed-rank tests with Bonferroni correction for non-normally distributed data. Error bars represent interquartile range for boxplots and P values below 0.05 were considered significant using the following notation: * p < 0.05, ** p < 0.01, *** p < 0.001, **** p < 0.0001.

To further examine differences in actin architecture between WT and PTPN22 KO Jurkat T cells at the mature synapse, we quantified the relative areas of actin coverage (Fig. 1F) and clearance (Fig. 1G), along with the number of F-actin-rich foci in the central region (Fig. 1H) at the 10 min timepoint. PTPN22 KO cells exhibited significantly reduced actin coverage, increased actin clearance, and a higher number of foci compared to WT. These quantitative characterizations support the observation of increased central actin concentration and reduced peripheral branching in PTPN22 KO cells relative to WT. The radial fluorescence intensity profiles (Fig. 1I), computed at 5- and 10-minutes post-activation, provides further comparisons of the dynamic cytoskeletal network rearrangements between WT and PTPN22 KO cells. At 5 minutes, WT cells exhibited minor actin clearance, whereas KO cells displayed features indicative of mature synapse formation at this early stage. By 10 minutes, the peripheral F-actin ring characteristic of a mature immunological synapse was observed in both cell types (r > 0.5). However, only PTPN22 KO cells exhibited a distinct central actin peak at the cSMAC, corresponding to F-actin foci structures.

### PTPN22 Modulates Arp2/3-Mediated Actin Remodeling in T Cell Synapse Formation

Differences in actin remodeling between activated WT and PTPN22 KO Jurkat T cells were characterized primarily by the presence of large concentrations of actin in the center of the cell, corresponding to actin foci mediated by the Arp 2/3 complex. To investigate whether PTPN22 influences Arp 2/3-mediated actin pathways in T cells, we conducted live-cell imaging experiments with the Arp 2/3 inhibitor CK666 (see Movie S3 and S4), which binds to, and stabilizes, the inactive form of the complex (*22*).

Figure 2A shows the temporal dynamics of actin remodeling in CK666-treated WT and PTPN22 KO Jurkat T cells interacting with aCD3/aCD28-coated surfaces. Overall, CK666-treated cells exhibited reduced spreading and prominent actin spikes surrounding the cell periphery compared to untreated cells. Notably, CK666 treatment decreased the percentage of WT cells forming synapses (90 ± 3 % to 63 ± 9 %), whereas PTPN22 KO cells remained unaffected, leading to a significantly higher synapse formation rate in PTPN22 KO compared to WT (Fig. 2B). Despite this, both WT and PTPN22 KO CK666-treated cells exhibited a significant reduction in spreading area, with no significant differences detectable between the two cell types (Fig. 2C). Importantly, despite the inhibition, dynamic actin movement and distinct actin structures remained observable in both cell types.

**Fig. 2.**
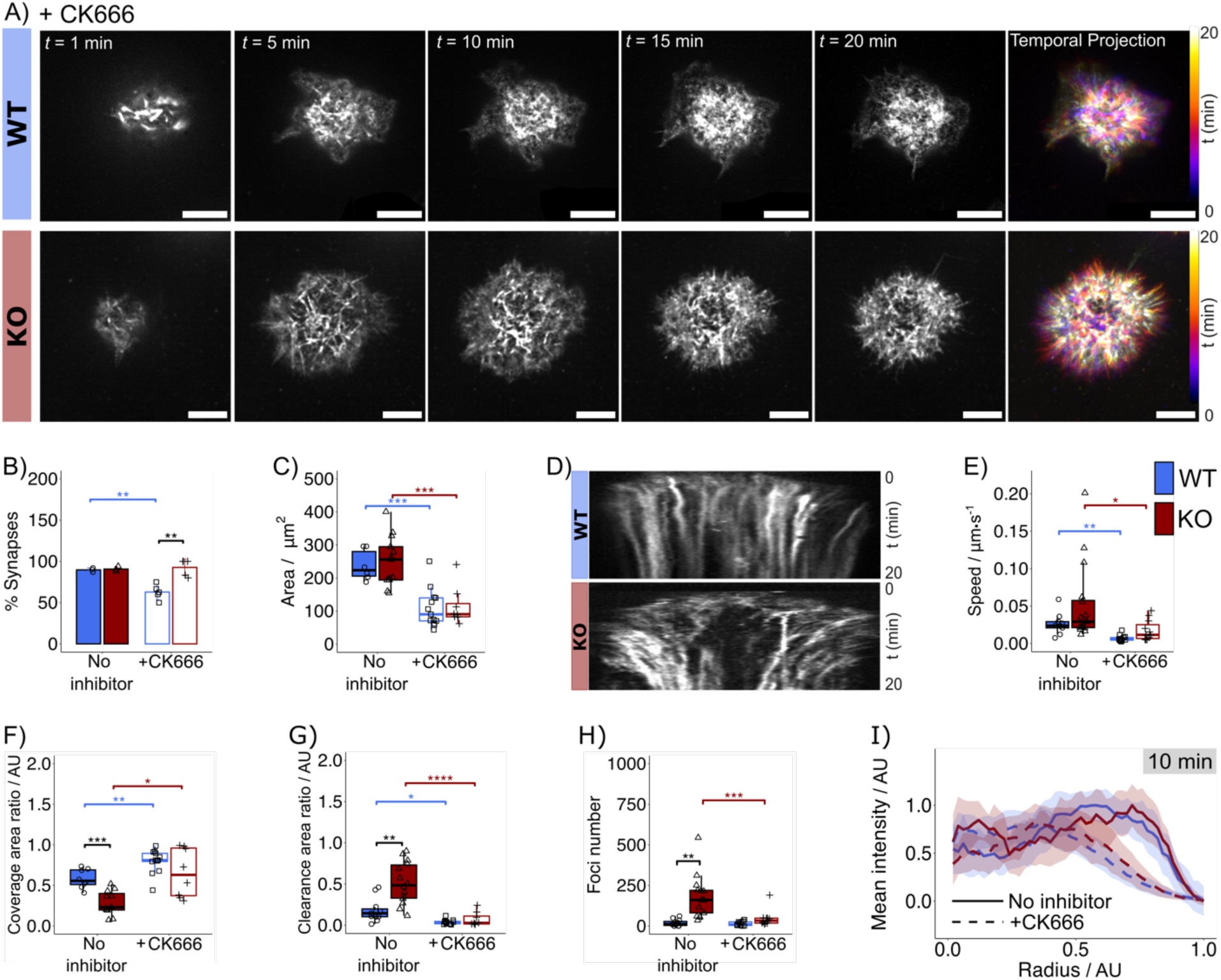
Actin remodeling in CK666 treated WT and PTPN22 KO Jurkat T cells interacting with aCD3/aCD28 coated slides. **(A)** Spinning disk confocal SoRa fluorescent imaging time- lapse of Jurkat T cells treated with the CK666 Arp 2/3 inhibitor and incubated with SiR actin from 1-20 minutes post settling. 1-, 5-, 10-, 15-, and 20-minute time frames are shown. Temporal projections of spreading are depicted with pseudo-color images with color gradients transitioning from purple to white representing early to late time-points. Scale bar = 5 μm. **(B)** The percentage of cells forming synapses on aCD3/aCD28 coated slides**. (C)** Maximum area of cell spreading. **(D)** Kymographs of CK666 treated aCD3/aCD28 activated WT and PTPN22 KO Jurkat T cells. **(E)** Initial speed of F-actin flow calculated from kymographs in **D**. **(F)** Ratio of F-actin ring coverage area to total cell area. **(G)** Ratio of central F-actin clearance area to total cell area. **(H)** The number of F-actin foci like structures found in the central regions of cells. **(I)** Radial analysis depicting mean relative F-actin intensity from the central region (radius = 0) to the edge of the cell (radius =1) after 10 minutes. Line indicates the mean with shaded areas representing standard deviation. For all plots except **B**, n = 10 cells from 3 independent passages. For **B**, n = 3 independent passages. Dashed lines and unfilled boxplots represent CK666 treated conditions with solid lines and filled boxplots representing untreated conditions. Statistical tests consisted of parametric ANOVA with pairwise Tukey’s HSD for normally distributed data and non-parametric Wilcoxon signed-rank tests with Bonferroni correction for non-normally distributed data. Error bars represent interquartile range for boxplots and P values below 0.05 were considered significant using the following notation: * p < 0.05, ** p < 0.01, *** p < 0.001, **** p < 0.0001.

CK666 treatment affects the formation of branched F-actin ring structures, which are crucial for T cell synapse formation and retrograde actin flow. Analysis of actin kymographs (Fig. 2D) indicated that CK666 treatment reduced initial retrograde actin flow speeds in both WT (0.7 ± 0.4 μm/min) and KO cells (1.8 ± 1.4 μm/min) compared to untreated cells (WT: (3 ± 2) x 10^−2^ μm/min; KO: (5 ± 5) x 10^−2^ μm/min) (Fig. 2E). Regarding actin architecture, CK666-treated WT cells exhibited increased linear actin features compared to untreated WT cells. In contrast, CK666-treated PTPN22 KO cells displayed a branched actin ring structure with central actin clearance by the mature synapse stage (15-20 minutes post-settling), differing from untreated PTPN22 KO cells, which exhibited a high central actin concentration and a diminished branched actin ring. Specifically, CK666-treated cells exhibited increased actin coverage (Fig. 2F) coupled with reduced actin clearance (Fig. 2G) compared to untreated cells. Notably, CK666-treated PTPN22 KO cells exhibited a significant reduction in central actin foci compared to untreated cells, suggesting a convergence in synaptic actin structures between WT and PTPN22 KO cells under CK666 treatment.

To further compare actin network rearrangements between CK666-treated and untreated WT and PTPN22 KO Jurkat T cells, radial fluorescence intensity profiles were computed at the 10-minute time point (Fig. 2I). CK666-treated cells exhibited central actin clearance characteristic of mature synapse formation, but at a radial distance closer to the cell center, suggesting a reduced cSMAC region. Strikingly, CK666-treated PTPN22 KO cells showed a substantial reduction in central actin, leading to an actin synapse structure more closely resembling that of WT cells under CK666 treatment. To confirm that these effects were specific to CK666 rather than the solvent DMSO, control experiments were conducted using DMSO-treated cells. Cells treated with DMSO alone and settled on aCD3/aCD28-coated slides exhibited phenotypes similar to untreated cells in both WT and PTPN22 KO Jurkat T cells (Fig. S2), indicating that DMSO did not alter actin dynamics. These findings suggest that PTPN22 primarily influences Arp 2/3-mediated actin remodeling, aligning with previous reports linking PTPN22’s binding partner PSTPIP-1 to WASp activation of the Arp 2/3 complex (*16*).

### PSTPIP-1 Redistribution and Cytoskeletal Interactions Modulate TCR Clustering in PTPN22 KO T Cells

PTPN22 has been identified as a direct binding partner of the cytoskeletal-associated protein PSTPIP-1 in T cells (*11*, *17*, *23*, *24*), suggesting that its potential role in actin remodeling may also be mediated through this interaction. However, PSTPIP-1 localization at the T cell synapse remains underexplored, and while its association with the tubulin cytoskeleton has been documented in monocytes (*25*), its distribution in T cells is less well understood. To characterize PSTPIP-1 localization and cytoskeletal interactions, Jurkat T cells (WT and PTPN22 KO) were activated on aCD3/aCD28 antibody-coated slides, fixed, and immunostained for PSTPIP-1, F-actin, and α-tubulin. Two polyclonal PSTPIP-1 antibodies were tested: (i) aPSTPIP-1 Ab1, targeting the PTPN22 binding region, and (ii) aPSTPIP-1 Ab2, targeting the CD2/WASp interaction region, to ensure unbiased detection. Antibody binding sites are illustrated in Fig. S3. PSTPIP-1 was visualized using AlexaFluor 568 Fab, F-actin with Alexa Fluor 405 Plus phalloidin, and α-tubulin with Alexa Fluor 488 conjugated antibody. Cells were imaged using the SoRa microscope with z-stacks of 100 nm steps up to 10 μm to capture 3D localization. Example images can be seen in Fig. S3A.

PSTPIP-1 staining using the two different antibodies revealed subtle differences in protein localization. aPSTPIP-1 Ab1 showed a stronger presence at the cell edges, whereas aPSTPIP-1 Ab2 was more diffusely distributed (Fig. S3A). Despite these differences, both antibodies clearly demonstrated clustering of PSTPIP-1 toward the center of activated cells, with Ab2 providing a more pronounced signal. Additionally, PTPN22 KO cells exhibited higher PSTPIP-1 staining compared to WT cells. The observed discrepancies may be due to differences in the antibody epitope binding regions. The binding of aPSTPIP-1 Ab1 may be hindered by PTPN22 in this region, potentially explaining why the increased central concentration and puncta of PSTPIP-1 were not visualized with this antibody in WT cells. In contrast, aPSTPIP-1 Ab2 binding could be influenced by CD2 and/or WASp binding. Since CD2 localizes to the peripheral pSMAC in activated T cells, this may account for why aPSTPIP-1 Ab2 staining did not highlight the peripheral PSTPIP-1 distribution as effectively as aPSTPIP-1 Ab1. To minimize potential bias from these differential epitope bindings, subsequent super-resolution fluorescence microscopy imaging used a combination of both antibodies.

Colocalization analysis revealed that PSTPIP-1 associates with both actin and tubulin in Jurkat T cells, in both unstimulated and activated conditions (Fig. S3B and S3C), as observed with both antibodies tested. Activation significantly increased colocalization, as shown by Pearson’s correlation coefficients, with no significant differences between WT and PTPN22 KO cells. This suggests that PTPN22 does not regulate PSTPIP-1’s cytoskeletal association. However, PSTPIP- 1 exhibited a higher central concentration in PTPN22 KO cells, where the TCR resides, making super-resolution fluorescence microscopy essential for gaining further insights into PSTPIP-1 clustering and its possible association with the TCR in the cSMAC region.

DNA-PAINT and Exchange-PAINT, combined with quantitative spatial analysis, were used to examine clustering and association of PSTPIP-1 and TCR in WT and PTPN22 KO Jurkat T cells (*26*, *27*). DNA-PAINT is a single-molecule localization microscopy technique that relies on the periodic binding of short DNA imager strands to docking strands, which are targeted to the protein of interest. Exchange-PAINT enables sequential DNA-PAINT imaging of multiple proteins through differential DNA strand binding. This approach provides high-resolution imaging of multiple proteins (with lateral resolutions on the order of 10 nm) and allows for precise protein quantification via qPAINT analysis (a schematic of this technique is provided in Fig. S4) (*26*, *28*). Cells were activated on aCD3/aCD28-coated slides, fixed, and stained with DNA-labeled antibodies targeting PSTPIP-1 and the TCRζ chain. DNA-PAINT imaging was performed using the SoRa microscope to leverage the large field of view (FOV) provided by spinning disk configuration (*29*). Sequential imaging with Cy3B-labelled complementary DNA strands for each target protein enabled their visualization at nanoscale resolution (median localization precision of 9 nm for both targets, Fig. S5).

Figure 3A displays representative DNA-PAINT images of TCR and PSTPIP-1 in WT and PTPN22 KO T cells activated with aCD3/aCD28 for 10 minutes, alongside SoRa fluorescence imaging of actin networks. Control experiments with T cells contacting bare glass surfaces are shown in Fig. S7. Visual inspection of activated T cells reveals a more pronounced central concentration of TCR proteins in PTPN22 KO cells compared to WT cells, with larger regions of high TCR coverage (Fig. 3A). This is accompanied by an increased presence of PSTPIP-1 in PTPN22 KO cells. Overlaying actin images with DNA-PAINT localizations of TCR and PSTPIP-1 localizations reveals their co-localization with actin foci, particularly in the central region of PTPN22 KO cells (Fig. 3A). To further analyze TCR and PSTPIP-1 spatial distribution, DNA-PAINT images were converted into protein maps (Fig. 3B and 3C) using the analysis routine detailed in Fig. S6. Protein density analysis across different cellular regions (center, mid, and edge; Fig. 3E) confirmed that TCR density was significantly higher in activated PTPN22 KO cells than in WT cells, while unstimulated cells showed no significant differences between WT and KO cells, with an average TCR density of approximately 28 proteins/μm² (Fig. S7E). PSTPIP-1 density, on the other hand, was significantly elevated in the mid and edge regions of unstimulated PTPN22 KO cells compared to WT cells (Fig. S7F). Upon activation, however, no significant differences in PSTPIP- 1 density were observed between WT and PTPN22 KO cells, despite a modest increase in the center and edge regions. Unlike TCR, which showed a significant increase in central protein density upon stimulation in both WT and KO cells, PSTPIP-1 density significantly decreased in activated PTPN22 KO cells. Despite this reduction, and following the trend observed for TCR, PSTPIP-1 density was significantly higher in the center of activated WT and PTPN22 KO cells compared to the mid region and was lower in the mid region compared to the edge (Fig. 3E).

**Fig. 3.**
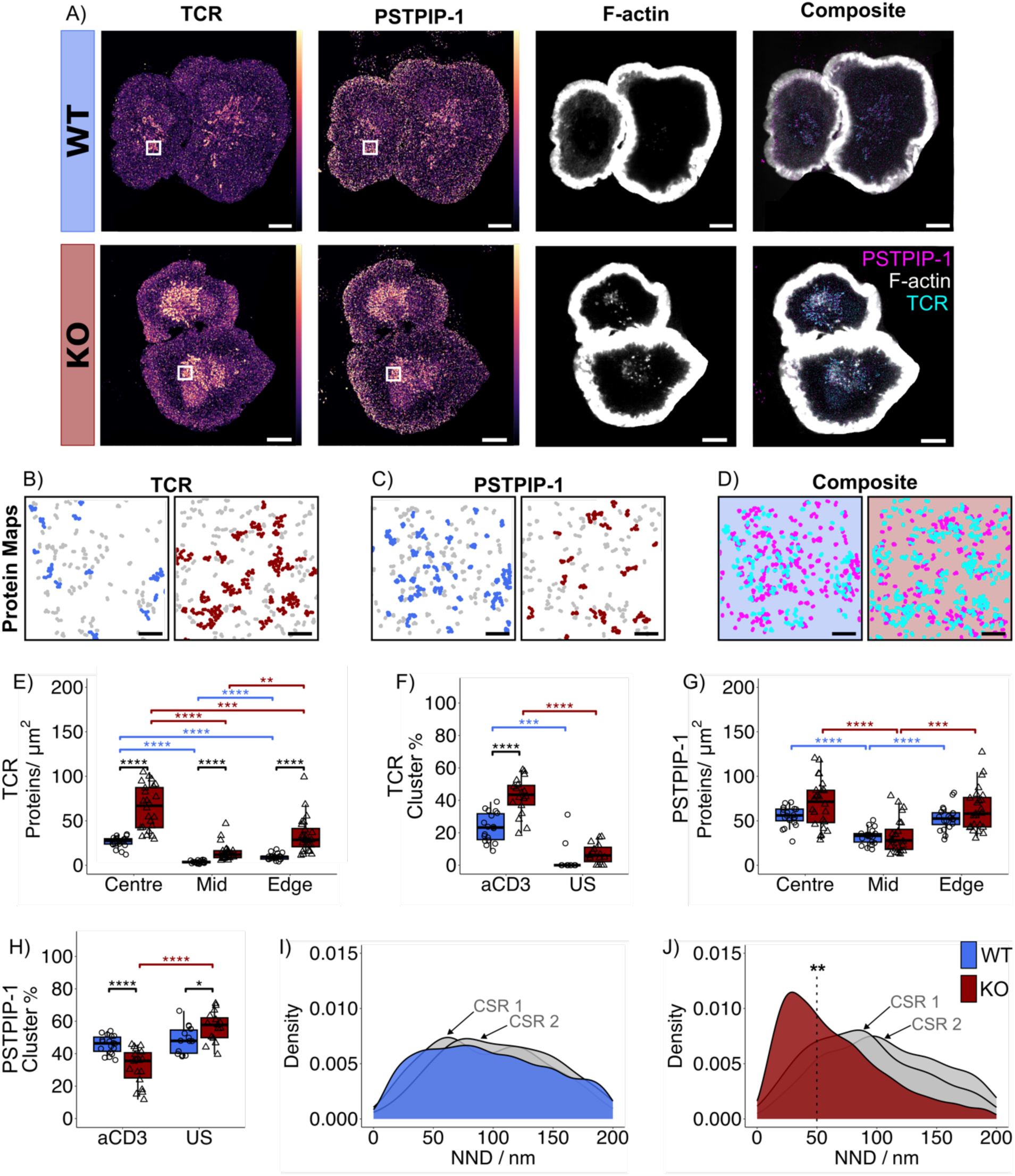
DNA-PAINT imaging and qPAINT quantification of TCR and PSTPIP-1 proteins coupled with SoRa super-resolution imaging of F-actin networks in aCD3/aCD28 activated WT and PTPN22 KO Jurkat T cells. **(A)** SoRa DNA-PAINT images of TCR and PSTPIP-1 coupled with fluorescent super-resolution images of F-actin in WT and PTPN22 KO Jurkat T cells after 10 minutes of activation via incubation on slides coated with aCD3/aCD28 antibodies. DNA- PAINT images colored by density with yellow to purple representing high to low density of single molecule localizations. Scale bar = 5 μm. **(B)** Zoom region of TCR protein maps calculated from the qPAINT pipeline as depicted by white boxes in **A**. Colored protein clusters contain 5 or more proteins. Scale bar = 500 nm. **(C)** Zoom region of PSTPIP-1 protein maps calculated from the qPAINT pipeline as depicted by white boxes in **A**. Colored protein clusters contain 5 or more proteins. Scale bar = 500 nm. **(D)** Composite overlay of TCR (cyan) and PSTPIP-1 (magenta) protein maps. Scale bar = 500 nm. **(E)** TCR protein density in 3 x 3 μm^2^ ROIs from the center, mid, and edge regions of aCD3/aCD28 activated cells as depicted in Fig. S7. **(F)** The percentage of TCR proteins in clusters of 5 proteins or more in central ROIs of all cells. **(G)** PSTPIP-1 protein density in 3 x 3 μm^2^ ROIs from the center, mid, and edge regions of aCD3/aCD28 activated cells. **(H)** The percentage of PSTPIP-1 proteins in clusters of 5 proteins or more in central ROIs of all cells. **(I)** Two-color 1^st^ nearest-neighbor distance (NND) analysis of PSTPIP-1 – TCR proteins in central ROIs of aCD3/aCD28 activated WT cells compared to complete spatial randomness models (CSR) shown in grey. **(J)** Dual-color 1^st^ NND analysis of PSTPIP-1 – TCR proteins in central ROIs of aCD3/aCD28 activated PTPN22 KO cells compared to CSR shown in grey. Dotted line depicts 50 nm range at which significance was tested. For all plots, n = 10 ROIs from 10 cells from 3 independent passages. Statistical tests consisted of parametric ANOVA with pairwise Tukey’s HSD for normally distributed data and non-parametric Wilcoxon signed-rank tests with Bonferroni correction for non-normally distributed data. Error bars represent interquartile range for boxplots and P values below 0.05 were considered significant using the following notation: * p < 0.05, ** p < 0.01, *** p < 0.001, **** p < 0.0001.

Quantification of TCR and PSTPIP-1 protein clustering is shown in Fig. S9, with the percentage of clustered proteins (defined as clusters of five or more proteins) presented in Fig. 3F for TCR and Fig. 3H for PSTPIP-1 (see Fig. S6-S9 for details on protein clustering analysis). Clustering analysis revealed a significant increase in TCR clustering at the center of both WT and PTPN22 KO cells upon activation. Notably, activated PTPN22 KO cells exhibited significantly higher TCR clustering than WT cells, consistent with the increased protein density. In contrast, while PSTPIP- 1 protein density did not differ significantly between activated WT and PTPN22 KO cells, clustering was greater in WT cells. In unstimulated cells, the opposite trend was observed, leading to a significant reduction in PSTPIP-1 clustering upon activation in PTPN22 KO cells. These findings highlight the importance of spatial organization in TCR and PSTPIP-1 signaling.

Dual-color Nearest Neighbor Distance (NND) analysis further examined the spatial relationship between PSTPIP-1 and TCR proteins at the center of the synapse (Fig. 3I, 3J). In activated PTPN22 KO cells, a significant association between TCR and PSTPIP-1 proteins was observed above density-matched complete spatial randomness (CSR) models (Fig. 3J), whereas no such association was detected in WT (Fig. 3I) or unstimulated cells (Fig. S7). The observed association between PSTPIP-1 and TCR in activated PTPN22 KO cells, suggests that in the absence of PTPN22, PSTPIP-1 localizes to the membrane, promoting actin foci formation and driving TCR clustering.

### PTPN22 Phosphatase Activity is Required for Regular Actin Remodeling in T Cells

Since the PSTPIP-1 binding region lies outside the phosphatase domain of PTPN22, it may be unlikely that PTPN22 directly dephosphorylates PSTPIP-1. However, its paralog, PTP-PEST (Protein Tyrosine Phosphatase - proline-, glutamic acid-, serine- and threonine-rich, also known as PTPN12), has been suggested to dephosphorylate PSTPIP-1 at specific tyrosine residues, such as Y344 (*30*). Yet, it is known that the phosphatase action of PTPN22 is affected by the autoimmune associated R620W mutation (*31*). This raises the question of whether PTPN22’s phosphatase activity is also required for appropriate regulation of actin remodeling in T cells or if PTPN22 only functions as a scaffolding protein, recruiting other actin-regulating factors, such as PSTPIP-1. To determine if PTPN22 phosphatase activity is required for actin remodeling, WT Jurkat T cells were pre-treated with the PTPN22-specific inhibitor LTV-1 (IC 50 of 508 nM) which competitively binds to the phosphate-binding loop of PTPN22, mimicking its natural substrate (*23*). As a readout, live-cell actin imaging was performed on cells layered onto aCD3/aCD28- coated slides (Movie S5).

Figure 4A presents the temporal evolution of actin remodeling in a representative WT cell treated with 5 μM LTV-1. LTV-1-treated WT cells exhibited altered actin remodeling compared to untreated WT cells (Fig. 1A), mimicking the PTPN22 KO phenotype. This alteration was characterized by actin accumulation in the central cSMAC area and depletion in the dSMAC and pSMAC regions, where the branched actin ring typically resides. Despite these structural changes, the overall synapse area remained similar between LTV-1-treated and untreated WT cells (Fig. 4B). To assess whether LTV-1-induced structural changes affected retrograde flow, kymographs were generated along the cell’s axis (Fig. 4C). Quantification of initial retrograde actin flow speeds (Fig. 4D) revealed no significant differences between LTV-1-treated WT cells, untreated WT cells, and PTPN22 KO cells. This suggests that while PTPN22 regulates actin remodeling, its phosphatase activity does not significantly influence retrograde actin flow.

**Fig. 4.**
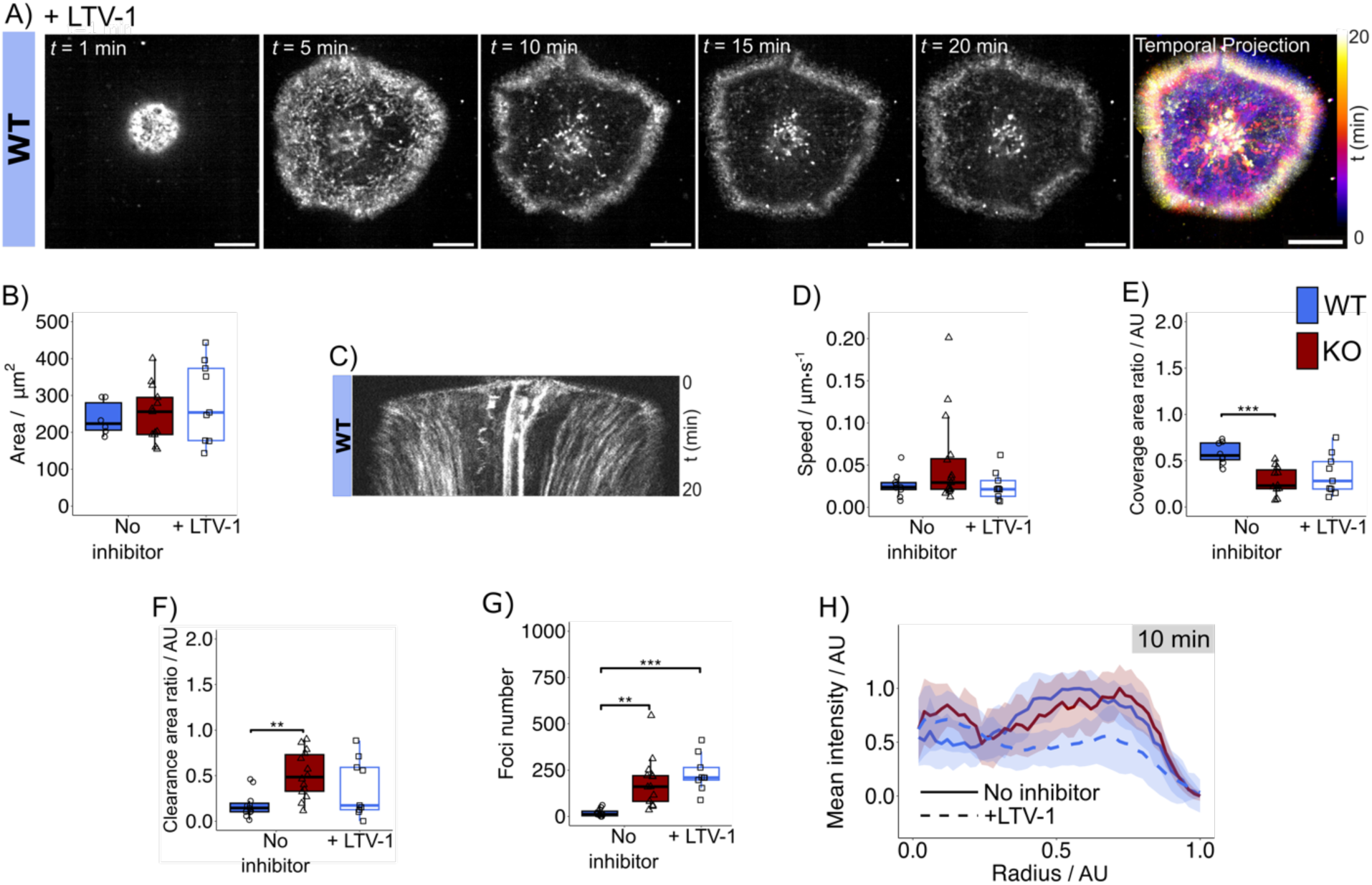
Actin remodeling in LTV-1 treated WT Jurkat T cells interacting with aCD3/aCD28 coated slides. **(A)** Spinning disk confocal SoRa fluorescent imaging time-lapse of Jurkat T cells treated with the LTV-1 PTPN22 phosphatase action inhibitor and incubated with SiR actin from 1-20 minutes post settling. 1-, 5-, 10-, 15-, and 20-minute time frames are shown. Temporal projections of spreading are depicted with pseudo-color images with color gradients transitioning from purple to white representing early to late time-points. Scale bar = 5 μm. **(B)** Maximum area of cell spreading. **(C)** Kymograph of an LTV-1 treated aCD3/aCD28 activated WT cell. **(D)** Initial speed of F-actin flow calculated from kymographs in **C**. **(E)** Ratio of F-actin ring coverage area to total cell area. **(F)** Ratio of central F-actin clearance area to total cell area. **(G)** The number of F- actin foci like structures found in the central regions of cells. **(H)** Radial analysis depicting mean relative F-actin intensity from the central region (radius = 0) to the edge of the cell (radius =1) after 10 minutes. Line indicates the mean with shaded areas representing standard deviation. For all plots, n =10 cells from 3 independent passages. Dashed lines and unfilled boxplots represent LTV-1 treated conditions with solid lines and filled boxplots representing untreated conditions. Statistical tests consisted of parametric ANOVA with pairwise Tukey’s HSD for normally distributed data and non-parametric Wilcoxon signed-rank tests with Bonferroni correction for non-normally distributed data. Error bars represent interquartile range for boxplots and P values below 0.05 were considered significant using the following notation: * p < 0.05, ** p < 0.01, *** p < 0.001, **** p < 0.0001.

To further examine actin architecture, the relative actin coverage and actin clearance areas were quantified (Fig. 4E, F). LTV-1-treated WT cells displayed a slight decrease in actin coverage and a slight increase in actin clearance compared to untreated WT cells, reflecting trends seen in PTPN22 KO cells, although these differences were not statistically significant. However, a significant increase in actin foci was observed in LTV-1-treated WT cells compared to untreated WT cells (Fig. 4G). This accumulation of actin foci at the cell center further aligns with the phenotype seen in PTPN22 KO cells, indicating that PTPN22 phosphatase activity plays a critical role in regulating actin foci formation.

Radial fluorescence intensity profiles were generated to compare actin distribution in LTV-1- treated WT cells, untreated WT cells, and PTPN22 KO cells (Fig. 4H). At the mature synapse stage (10 minutes post-settling), LTV-1-treated WT cells displayed increased central actin accumulation and a well-defined actin ring at the cell periphery (r > 0.5), mirroring the spatial distribution observed in PTPN22 KO cells. In contrast, untreated WT cells exhibited central actin clearance, characteristic of a mature synapse. These results further confirm the fact that PTPN22 phosphatase activity is required for proper T cell actin remodeling.

### Actin Remodeling and Calcium Signaling are Significantly Altered in PTPN22 KO T Cells upon Low-Affinity Activation

Both PSTPIP-1 and PTPN22 are T cell proteins known to harbor mutations associated with autoimmune and autoinflammatory disorders. Autoimmunity is inherently linked to immune tolerance mechanisms normally associated with low affinity TCR responses to self, which when breeched leads to inappropriate immune reactions. To better understand how PTPN22 contributes to these processes, we utilized WT and PTPN22 KO TCR⁻/⁻ Jurkat T cells engineered to express a transgenic TCR with high affinity for the pTax peptide and low affinity for the pHuD peptide. These engineered cells provide a controlled system to investigate how PTPN22 influences actin remodeling and calcium signaling in response to TCR-peptide-MHC (Major Histocompatibility Complex) interactions of differing affinities. To visualize these processes, cells were labeled with SiR-actin dye and imaged on slides coated with either high-affinity (pTax) or low-affinity (pHuD) ligands displayed on the MHC-like molecule Dimer XI (Movies S6- S9).

Fig. 5A displays the temporal evolution of actin remodeling in WT and PTPN22 KO cells settling on pTax-coated slides, notably these cells displayed less pronounced differences in actin architecture. In PTPN22 KO cells, peripheral branched actin was present, but central actin concentration was lower compared to cells stimulated with aCD3/aCD28 (Fig. 1A). WT cells retained the characteristic three-zone F-actin structure, though these zones appeared later and were less sharply defined than in aCD3/aCD28-stimulated cells. Strikingly, under low-affinity conditions, PTPN22 KO cells exhibited an exceptionally dense concentration of actin in the central region, with less defined peripheral actin structures when compared to WT cells which showed only a modest increase in central actin concentration (Fig. 5B). This suggests that PTPN22 plays a key role in regulating actin dynamics in response to weak TCR signaling.

**Fig. 5.**
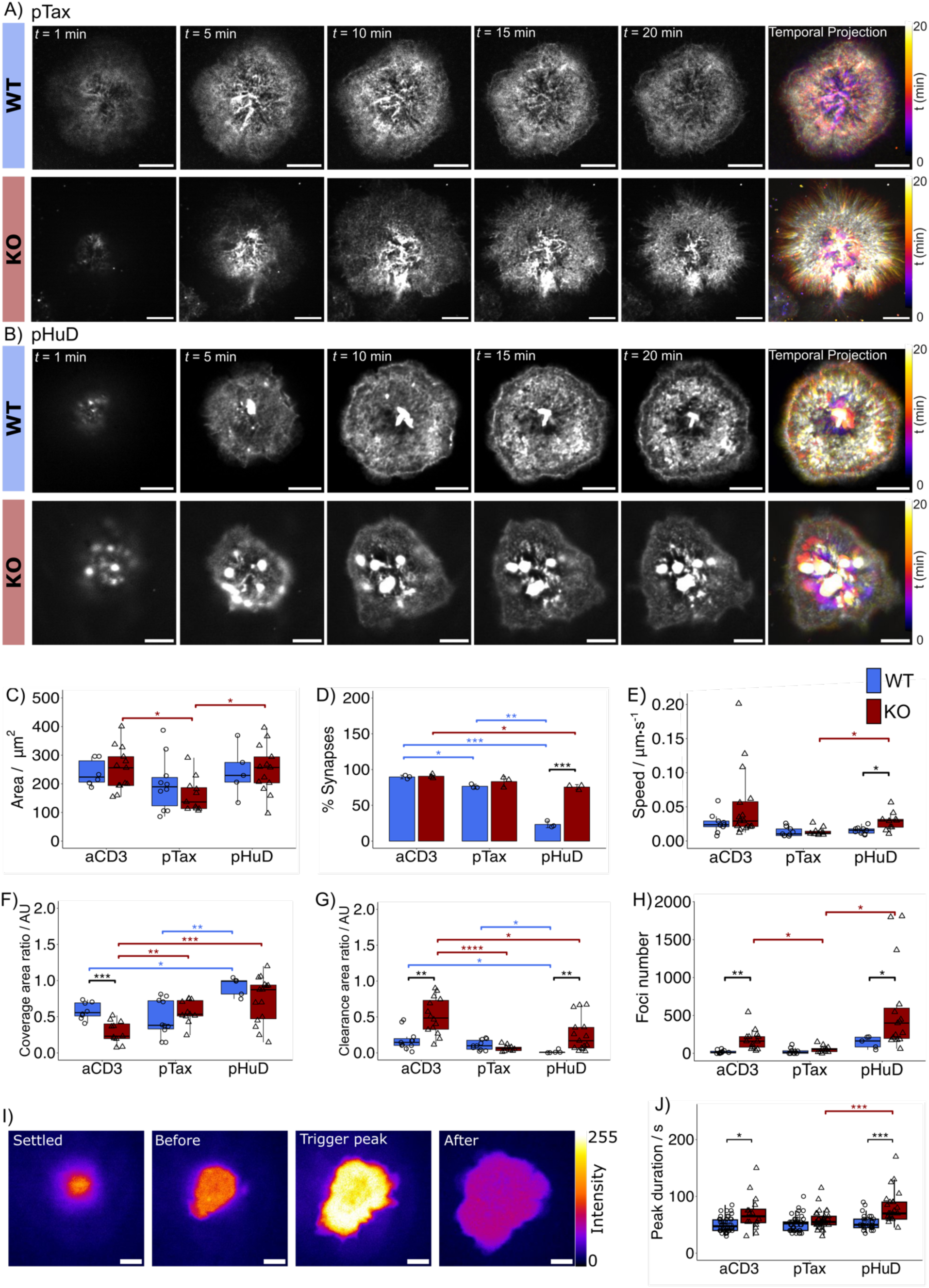
Actin remodeling and calcium flux in WT and PTPN22 KO Jurkat T cells interacting with high and low affinity pMHC coated slides. (A-B) Spinning disk confocal SoRa fluorescent imaging time-lapse of Jurkat T cells incubated with SiR actin from 1-20 minutes post settling. 1-, 5-, 10-, 15-, and 20-minute time frames are shown. Temporal projections of spreading are depicted with pseudo-color images with color gradients transitioning from purple to white representing early to late time-points. High affinity pTax pMHC images are shown in **A** with low affinity pHuD pMHC images shown in **B**. Scale bar = 5 μm. **(C)** The percentage of cells forming synapses on aCD3/aCD28, high affinity pTax, and low affinity pHuD coated slides. **(D)** Maximum area of cell spreading. **(E)** Initial speed of F-actin flow. **(F)** Ratio of F-actin ring coverage area to total cell area. **(G)** Ratio of central F-actin clearance area to total cell area. **(H)** The number of F-actin foci like structures found in the central regions of cells. **(I)** Example SoRa fluorescent images of a cell labelled with Calbryte 520 calcium ion dye experiencing a calcium burst over time, colored by intensity, scale bar = 5 μm. **(J)** The duration of the peak calcium signal in transient calcium signaling cells. For all plots, n =10 cells from 3 independent passages. Statistical tests consisted of parametric ANOVA with pairwise Tukey’s HSD for normally distributed data and non-parametric Wilcoxon signed-rank tests with Bonferroni correction for non-normally distributed data. Error bars represent interquartile range for boxplots and P values below 0.05 were considered significant using the following notation: * p < 0.05, ** p < 0.01, *** p < 0.001, **** p < 0.0001.

The ability of cells to form synapses was highly dependent on ligand affinity (Fig. 5C). WT cells formed synapses most efficiently on aCD3/aCD28-coated surfaces (90 ± 3%), followed by pTax- coated surfaces (77 ± 3%). In contrast, synapse formation dropped significantly under low-affinity pHuD conditions (23 ± 5%). However, PTPN22 KO cells exhibited a strikingly different pattern, forming synapses at similar rates across all ligand conditions, including a significantly increased percentage under pHuD activation compared to WT. This suggests that PTPN22 KO cells have reduced sensitivity to ligand affinity during synapse formation.

While the final contact area between WT and PTPN22 KO cells remained comparable across conditions (Fig. 5D), PTPN22 KO cells formed larger contact areas under aCD3/aCD28 and pHuD activation when compared to the high-affinity pTax activation. Analysis of actin flow also revealed significant differences for PTPN22 KO cells activated under low-affinity conditions. In WT cells, F-actin exhibited inward movement at a speed of (3 ± 1) × 10^−2^ μm/min under aCD3/aCD28 activation, with similar flow rates observed for pTax (1.4 ± 0.7 × 10^−2^ μm/min) and pHuD (1.6 ± 0.5 × 10^−2^ μm/min) conditions (Figure 5E). PTPN22 KO cells, however, displayed significantly faster actin flow under low-affinity pHuD activation compared to WT cells and the activation under high-affinity pTax for PTPN22 KO cells, further supporting the idea that PTPN22 regulates actin remodeling in a ligand-affinity-dependent manner.

Regarding actin architecture, WT cells exhibited significantly increased F-actin coverage (Fig. 5F) and reduced central actin clearance (Fig. 5G) under low-affinity pHuD conditions, compared to aCD3/aCD28 or pTax conditions. Additionally, WT cells showed a slight, though not statistically significant, increase in F-actin foci under these conditions (Fig. 5H). Conversely, PTPN22 KO cells activated under high-affinity pTax conditions displayed significantly greater actin coverage (Fig. 5F), reduced central actin clearance (Fig. 5G), and fewer foci (Fig. 5H) compared to aCD3/aCD28 conditions. Similar trends were observed under low-affinity pHuD activation, where PTPN22 KO cells exhibited higher actin coverage (Fig. 5F) and reduced clearance (Fig. 5G) but showed no significant difference in foci number compared to aCD3/aCD28 activation conditions. This suggests that loss of PTPN22 disrupts actin clearance and promotes excessive central actin accumulation, particularly under weak TCR signaling.

To examine whether PTPN22 KO cells exhibit altered calcium flux upon stimulation, live-cell calcium imaging was performed using the Calbryte 520 dye on the SoRa fluorescence microscope capturing an image every 5 seconds over a 20-minute period (Fig. 5I). T cells were tracked from initial settling on ligand-coated surfaces to peak calcium ion release, capturing the early signaling events that coincide with actin remodeling. WT cells exhibited similar calcium flux durations across all activation conditions (Fig. 5J). However, PTPN22 KO cells showed significantly prolonged calcium elevation peaks under both aCD3/aCD28 and low-affinity pHuD conditions. Additionally, PTPN22 KO cells displayed longer periods of calcium flux under pHuD compared to high-affinity pTax activation. This suggests that in the absence of PTPN22, weak TCR signaling leads to prolonged calcium signaling, potentially contributing to sustained downstream activation phenotypes previously reported in the literature (*9*, *20*, *32*).

## DISCUSSION

In this study, we investigated the role of PTPN22 in actin remodeling during T cell synapse formation, highlighting its regulation of actin pathways under conditions of different ligand binding affinity. Our findings uncover a role for PTPN22 phosphatase activity in modulating actin dynamics, particularly in Arp2/3 complex-mediated remodeling and central F-actin foci formation. Specifically, the loss of PTPN22 or its phosphatase activity leads to excessive central synaptic F-actin structures. We propose that this phenomenon may be driven by the prolonged phosphorylation, and thus activation, of a PSTPIP-1-associated actin-binding protein, with WASp being a likely candidate. WASp is directly regulated by PSTPIP-1, and its phosphorylation at tyrosine 291 by Fyn promotes Arp2/3-mediated actin polymerization(*33*, *34*). Previous studies have demonstrated that WASp is downregulated by PTP-PEST, another phosphatase in the same family as PTPN22 (*33*). Given these interactions, it is plausible that the absence of PTPN22 results in WASp overstimulation, leading to excessive Arp2/3 activation and the formation of F-actin foci at the expense of other synaptic actin structures, such as the branched actin ring (*35*).

Further supporting our model that PTPN22-dependent aberrant actin remodeling occurs through its interaction with PSTPIP-1 is a recent study on patient samples carrying the R228C mutation, which disrupts the interaction between PSTPIP-1 and PTPN22 (*17*). In this study, a greater decrease in F-actin content was observed in CD4+ T cells from patients carrying the mutation compared to healthy donors after anti-CD3 stimulation (*18*). These findings align with our observations of increased actin clearance in PTPN22 KO cells, reinforcing the idea that PTPN22 regulates synaptic actin dynamics by modulating PSTPIP1-associated pathways.

Beyond its role in actin remodeling, our results show that PTPN22 also appears to influence TCR clustering. The increased density of TCRs and TCR clusters in the cSMAC region of PTPN22 KO cells suggests that actin remodeling may regulate receptor distribution at the synapse. One possible mechanism for this observation is that the excess F-actin foci could stabilize TCR microclusters, prolonging their residence time at the synapse. Additionally, actin remodeling is known to play a crucial role in receptor endocytosis and recycling. In T cells, this process ensures the continuous recycling of surface receptors, including the TCR, which is essential for both sustained signaling and eventual signal termination. The WASH complex, which promotes Arp2/3-mediated actin polymerization, is critical for receptor retrieval and recycling. Previous studies have demonstrated that loss of WASH results in reduced TCR surface clustering due to impaired recycling (*35*). Given that PTPN22 KO cells display excessive actin foci and increased TCR clustering, it is possible that PTPN22 deficiency also leads to overactivation of WASH, shifting receptor trafficking toward recycling rather than degradation. This hypothesis is supported by recent findings showing that reduced PTPN22 phosphatase activity promotes CD3ε recycling over internalization (*36*).

The increased TCR clustering in PTPN22 KO cells has significant functional consequences, particularly in enhancing T cell activation. Larger TCR clusters are associated with greater signaling potency, as they facilitate the accumulation of key signaling molecules such as CD3ζ ITAMs (Immune Tyrosine Activation Motifs), ZAP-70 (Zeta-chain-Associated Protein kinase 70), and LAT (Linker of Activated T cells) (*37*). Our results demonstrate that PTPN22 KO cells exhibit prolonged calcium signaling, consistent with an increase in TCR stability. This prolonged calcium flux may result from enhanced PLC-γ phosphorylation, leading to sustained NFAT nuclear localization and increased cytokine production (*38*). Notably, these effects were most pronounced under conditions of low-affinity TCR engagement. In WT cells, low-affinity TCR-pMHC interactions typically do not generate sufficient signaling-competent clusters to initiate activation (*39*). However, PTPN22 KO cells formed synapses at similar levels regardless of ligand affinity, indicating a loss of affinity discrimination. This suggests that PTPN22 plays a crucial role in modulating actin remodeling and TCR clustering in response to ligand affinity, thereby maintaining proper activation thresholds. This is in line with a growing body of literature highlighting the critical activity of PTPN22 in T cell activation to weak affinity stimulation and loss of affinity sensitivity (*20*, *40–42*). PTPN22 has also been highlighted as a possible target for cancer immunotherapy, given the nature of increased T cell signaling upon PTPN22 KO and response to weak affinity ligands (*41*, *43–45*).

Finally, our study describes the localization of PSTPIP-1 in both unstimulated and stimulated T cells, as well as its colocalization with TCRs in the synapse. In unstimulated cells, PSTPIP-1 is distributed across both actin and microtubule structures, but upon T cell activation, it becomes more concentrated within central actin foci. This redistribution suggests a dynamic role for PSTPIP-1 in coordinating cytoskeletal changes upon activation. Interestingly, Burn et al. (*46*) also shown similar localization and clustering behavior of PTPN22 in unstimulated WT T cells and cells undergoing ICAM-1-LFA-1 engagement. Moreover, the increased colocalization of PSTPIP- 1 with TCRs in PTPN22 KO cells further supports its involvement in receptor clustering and recycling. Given its interactions with both actin regulators and signaling molecules, PSTPIP-1 may serve as a key scaffold for organizing signaling microdomains at the synapse.

In summary, our findings demonstrate that PTPN22 plays a crucial role in regulating actin remodeling, TCR clustering, and signaling dynamics at the T cell synapse. The absence of PTPN22 leads to exaggerated actin foci formation, increased TCR clustering, and prolonged calcium signaling, particularly under low-affinity conditions. These observations underscore the importance of PTPN22 in maintaining proper T cell activation thresholds and suggest potential implications for autoimmune disease mechanisms, where loss of activation control may contribute to aberrant immune responses.

## MATERIALS AND METHODS

### Cell culture

WT and PTPN22 KO TCR⁻/⁻ Jurkat T cells, engineered to express a transgenic TCR with high affinity for the pTax peptide (LLFGYPVYV, from human T-lymphotropic virus type 1; KD = 1.8 μM) and low affinity for the pHuD peptide (LGYGFVNYI, from human neuronal protein; KD = 123 μM), were obtained from Prof. Rose Zamoyska at the University of Edinburgh and used as the primary samples for this study. This TCR system, known as the Tax TCR, was modified by CRISPR knockout of PTPN22 Exon 1 to generate the PTPN22 KO cells, as described in the publication by Bray et al. (*20*). Western blot and PTPN22 Exon 1 region sequencing confirmed the deletion of PTPN22 protein in the KO cell line. Cell lines were cultured in Roswell Park Memorial Institute (RPMI) media (Thermo fisher), supplemented with 10 % fetal bovine serum (FBS, Gibco) and 1 % penicillin with streptomycin (P/S, Gibco). Cells were maintained at 1 x 10^5^ cells/mL and cultured at 37 °C and 5 % CO2.

### Cell activation

WT and PTPN22 KO Jurkat T cells were activated on glass bottom 6 channel Ibidi µ-slides (IB- 80607) functionalized with activating ligands of differing affinities. For antibody activation 50 μL of 2 μg/mL anti-CD3 (OKT3, Invitrogen) and 5 μg/mL anti-CD28 (CD28.2, Invitrogen) in phosphate buffered saline (PBS, LifeTech) was added to chambers and incubated overnight at 4 °C. For peptide stimulation, pTax and pHuD peptides (GenScript) were reconstituted in Dimethyl sulfoxide (DMSO, Thermo Fisher) at 5 mg/mL. High-affinity activation involved incubating 10 μg/mL of dimeric human HLA-A2:Ig fusion protein, known as Dimer XI, (BD Biosciences) with a 160× molar excess of pTax peptide (2 mg/mL) in PBS overnight at 37 °C and 5 % CO2. Low-affinity activation used the same method with a 640× molar excess of pHuD peptide. After incubation, 50 μL was added to the chambers and incubated overnight at 4°C.

For live cell imaging, unstimulated controls were placed on bare glass slides washed with 100 μL of filtered PBS. For fixed cell imaging, slides were functionalized with 0.01% poly-L-lysine (PLL, Sigma-Aldrich) via 15-minute incubation at room temperature (RT) to prevent cell detachment. All slides were washed with PBS through two 3-minute incubations after ligand treatment.

### Live cell imaging

For actin imaging, 5 x 10^5^ cells were incubated overnight at 37 °C and 5 % CO2 in 1 mL media containing 200 nM Silicon Rhodamine (SiR) actin live-cell dye (Spirochrome). For calcium ion imaging, 5 x 10^5^ cells were incubated for 1 hour at 37 °C and 5 % CO2 in 1 mL media containing 1 μM Calbryte 520 acetoxymethyl ester dye (AAT Bioquest), washed, centrifuged (140 x g, 7 minutes), and resuspended in 1x live-cell imaging solution (Invitrogen) at 1 x 10^6^ cells/mL. Slides functionalized with ligands were incubated with 100 μL of 1 x 90 nm gold nanoparticles (Cytodiagnostics) for 15 minutes, followed by PBS washing (2 x 3-minute incubations).

Live cell imaging was performed on a dual-camera (ORCA-Fusion BT Digital CMOS) Spinning Disk Super Resolution by Optical Pixel Reassignment (SoRa) microscope (CSU-W1 SoRA, Nikon) using a 60x objective (1.49 NA) at 37 °C and 5% CO2. Firstly, the microscope slide was placed on the sample mount for 10 minutes to warm to 37 °C. Gold nanoparticles were used to focus on the slide plane with a 638 nm laser at 70% power. Once the focus was found, 100 μL of cell solution was added to a chamber and imaging was immediately started to capture cells settling and activating on the ligand-coated glass slide.

For actin imaging, cells were imaged every 10 seconds for 20 minutes using a 638 nm laser at 30% power and a 708/75 bandpass filter. For calcium imaging, cells were imaged every 5 seconds for 20 minutes with a 488 nm laser at 20% power and a 525/50 bandpass filter. 2.8x and 1x SoRA magnification was used for actin and calcium imaging, respectively, rendering a pixel size of 39 nm and 108 nm.

For Arp 2/3 inhibition, 25 μM CK666 (Sigma-Aldrich) was added to cell suspensions for 3 minutes before imaging. For PTPN22 phosphatase activity inhibition, 5 μM LTV-1 (Sigma-Aldrich) was added 45 minutes before imaging. DMSO controls in which cells received the same volume of the solvent as in the treatments were also conducted.

After acquisition, all images were deconvoluted using the Richardson-Lucy 2D method with a maximum of 20 iterations and an effective pinhole size of 0.01 in the Nikon NIS-Elements software.

### Antibody – DNA coupling for DNA-PAINT

DNA docking strands were coupled to antibodies for DNA-PAINT imaging via a maleimide- PEG2-succinimidyl ester coupling reaction (*9*). Briefly, 13 μL of a thiolated DNA docking strand (Eurofin Genomics) was reduced with 30 μL of 250 mM dithiothreitol (DTT, LifeTech) for 2 hours. 30 minutes after starting this reaction, 175 μL of 0.8 mg/mL antibody solution was incubated with 0.9 μL of 23.5 mM maleimide-PEG2-succinimidyl ester cross-linker solution (Sigma-Aldrich) at 4 °C in the dark for 90 minutes. Both reactions were then purified via Microspin Illustra G-25 columns (GE Healthcare) and Zeba spin desalting columns (7K MWCO, Thermo Fisher Scientific), respectively. The DNA docking strand and antibody-linker solution were then mixed and incubated overnight at 4°C. Amicon spin filtration (Merck/EMD Millipore) was then performed to remove any uncoupled DNA.

A NanoDrop One spectrophotometer (Thermo Fisher Scientific) was used to measure DNA-antibody concentration and coupling ratio, yielding approximately 1:1 in all conjugation reactions. The antibodies and their coupled docking sequences were anti-rabbit-Fab antibody (Jackson ImmunoResearch), coupled to the docking strand 7xR4 (CTCTCTCTCTCTCTCTCTC), to target PSTPIP-1 primary antibodies and the anti-TCRζ antibody (6B10.2, BioLegend), coupled to the docking strand 7xR3 (ACACACACACACACACACA), for TCR.

### Fixed immunofluorescence and DNA-PAINT imaging

Cells were incubated at 5 x 10^5^ cells/mL in fresh media for 24 hours, centrifuged at 140 x g for 7 minutes, and resuspended in PBS at 1 x 10^6^ cells/mL. After ligand coating, slides were incubated at 37 °C and 5% CO2 for 15 minutes. 100 μL of cell suspension was added and incubated for 10 minutes (aCD3/aCD28 activation). Cells were fixed with 4% paraformaldehyde (PFA, LifeTech) for 30 minutes at room temperature (RT), followed by washing with 60 mM glycine (MP Biomedicals) in PBS to quench autofluorescence. Permeabilization was done with 0.1% Triton X- 100 (VWR) for 5 minutes, and samples were blocked with 5% Bovine Serum Albumin (BSA, Merck) in PBS for 60 minutes at RT.

For immunofluorescence, anti-PSTPIP-1 (Ab1 – polyclonal, Abcam, Ab2 – polyclonal, Novus Biologicals) and anti-α-Tubulin AF488 (DM1A, Abcam) were used at 5 μg/mL, and incubated for 1 hour at 37 °C. For DNA-PAINT, anti-PSTPIP-1 and DNA-coupled anti-TCRζ antibodies were used at 5 μg/mL and 10 μg/mL, respectively, for 1 hour at 37°C. Subsequent incubations were in the dark. Slides were washed, then secondary antibodies (anti-rabbit Fab AF568, Thermo Fisher for immunofluorescence, DNA-coupled anti-rabbit Fab for DNA-PAINT) were applied at 2 μg/mL for 1 hour at RT. F-actin was stained with AF405 plus phalloidin (Thermo Fisher) for 15 minutes at RT.

For DNA-PAINT, slides were incubated with 90 nm gold nanoparticles for 15 minutes, washed, and then imaged using a DNA imager solution containing 2 nM DNA imager, 1 mM ethylenediaminetetraacetic acid (EDTA, LifeTech), and 500 mM NaCl (Merck) in PBS.

Immunofluorescence imaging was performed on a SoRa microscope with a 60x objective lens (NA = 1.49), 2.8x SoRa magnification and 2x binning (pixel size of 78 nm). Z stacks were acquired with a 100 ms integration time and 0.1 μm axial step size. Sequential imaging was done using a 561 nm laser at 50% intensity (PSTPIP-1), a 488 nm laser at 20% intensity (α-Tubulin), and a 405 nm laser at 30% intensity (F-actin), with appropriate filters (605/52, 525/50, 447/60 bandpass).

For Exchange-PAINT imaging(*27*), slides were equipped with elbow Luer connector male adaptors and 0.5 mm silicon tubing (Ibidi) for fluid exchange. DNA-PAINT images were acquired on a SoRa microscope with a 60x objective lens (NA = 1.49) at 2.8x SoRa magnification and 2x2 binning, resulting in a pixel size of 78 nm. Imaging was performed with a 200 ms integration time, capturing 10,000 frames per round at 100% power using a 561 nm laser and a 605/52 band-pass filter. PSTPIP-1 and TCRζ were imaged sequentially. To image TCRζ, the complementary DNA imager solution to the DNA-coupled TCRζ antibody (TGTGTGT, Eurofin Genomics) was added, followed by washing with 10 mL PBS containing 1 mM EDTA and 500 mM NaCl to ensure complete removal of the first imager strand. The complementary DNA imager solution for PSTPIP-1 (GAGAGAG, Eurofin Genomics) was then added to image PSTPIP-1. Both imager strands were tagged with Cy3b dye at the 3’ end for imaging. A schematic of the DNA-PAINT imaging process is provided in Fig. S4.

### Live cell actin image analysis Percentage of cells forming synapses

The percentage of cells forming synapses was calculated by identifying cells that formed a contact area with the glass slide exceeding 100 μm^2^ and displayed an actin ring structure. For untreated and LTV-1-treated cells, synapse formation was assessed after 20 minutes of activation. For CK666-treated cells, synapses were defined by the presence of contact with the slide, an actin ring, and visible actin flow within the 20-minute imaging period. The percentage was calculated by the ratio of the number of synapse-forming cells to the total cells per field of view (FOV).

### Radial analysis

Radial analysis was performed by defining a circle around each cell in live-cell images at specific time points. Using the Fiji plugin “Radial Profile plot”, the intensity at increasing radii from the cell center to edge was measured (*47*). Min-max scaling was applied to obtain relative radius and intensity values for each cell.

### Area analyses

To obtain a mask of the cell contact area, images were Gaussian blurred with a 0.5 μm radius, then converted to a mask using the Fiji “Convert to Mask” function with the IsoData method. The total cell area in each frame was calculated using Fiji’s “Analyze Particles” function (10-500 μm^2^ area, including holes), and frame times were multiplied by 10 to give the area over time in seconds.

For actin coverage, the same procedure was repeated without Gaussian blurring and excluding holes in the “Analyze Particles” function. The ratio of actin area to total cell area was calculated by dividing the actin area by the total cell area.

To calculate actin clearance in the cell center, the actin area mask was inverted using Fiji’s “Binary Invert” function. Connected component labeling was performed using the napari software “napari-segment-blobs-and-things-with-membranes” package (*48*). The “Regionprops” function in napari was used to measure the area of each connected component. The area of actin clearance in the center was selected from the label list and divided by the total cell area to determine the clearance area ratio.

### Foci quantification

Foci were quantified using the following equation:

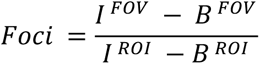

where I^FOV^ is the raw integrated density of the FOV covering the actin clearance area with foci; B^FOV^ is the raw integrated density of the FOV background; I^ROI^ is the raw integrated density of the selected 0.1 μm region of interest (ROI) corresponding to the smallest foci; B^ROI^ is the raw integrated density of the 0.1 μm background ROI. The background density, B^FOV^ was calculated from B^ROI^ using the following equation:

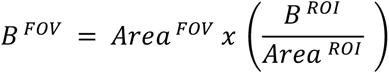

All values were measured in Fiji.

### Kymograph analysis

Kymographs were generated using Fiji’s “Reslice” function along a one-pixel thick line through the center of the cell. The speed of movement was calculated by dividing the change in distance by the change in time, using the xy coordinates of selected filaments from the Fiji segmented line tool. Segments were taken 1 μm from the cell edge, covering the mesh ring area.

### Live cell calcium image analysis

Calbyte 520 dye intensity was measured in 5 x 5 μm^2^ ROIs for each cell using Fiji, with background correction. Cells that experienced a calcium burst (triggering cells) were identified by a threshold of 0.2 times the maximum Calbyte intensity (min-max scaled) during the imaging period. Triggering cells were categorized into three types: transient (one short burst), oscillatory (multiple short bursts), and sustained (a burst followed by a sustained high signal). To define these categories min-max scaling was applied per cell, and the mean intensity was calculated. Cells with a mean intensity above 0.4 were categorized as sustained, while those below 0.4 with one peak were transient, and those with two or more peaks were oscillatory. For transient cells, peak duration was calculated as the length of time above a mean intensity of 0.5 for a single peak. All analyses were conducted in R (*49*).

### Fixed image analyses

After acquisition, 3D deconvolution was performed using the Richardson-Lucy method in Nikon Elements software with a pinhole size of 0.01 and 20 iterations. An axial step of 0.1 μm was used to achieve oversampling, improving axial resolution by up to 0.1 μm, as the Z-plane resolution of the SoRa microscope is 0.3 μm.

Z axis profile: Intensity in the Z plane of PSTPIP-1 3D Z-stack images was measured using the “Plot Z Axis Profile” function in Fiji on background corrected images.

Colocalization analysis: Colocalization of PSTPIP-1 with F-actin and α-tubulin structures in 3D Z-stack images was calculated using the Fiji plugin JACoP, with Pearson’s correlation coefficient determined on thresholded images (IsoData method) on a cell-by-cell basis. Actin and tubulin images were combined using the “add” function in Fiji.

### DNA-PAINT and qPAINT image analyses

After image acquisition, DNA-PAINT images were deconvoluted using the Blind method in Nikon Elements with a pinhole size of 0.01 and 12 iterations. The images were then processed in Picasso software (*26*), where single molecule blinking events were identified and fit with a 2D Gaussian function to create localization maps. Drift correction was performed using gold nanoparticles as fiducial markers, followed by additional drift correction via the redundant cross-correlation (RCC) method. Localization errors were filtered to 15 nm.

PSTPIP-1 and TCRζ localization maps were aligned using gold nanoparticles. Protein quantification was performed by analyzing the time between blinking events (dark time, τd), which decreases as the number of docking sites (proteins) increases. To conduct qPAINT analysis, single molecule localization maps were subject to cluster analysis. Density-Based Spatial Clustering of Applications with Noise (DBSCAN) algorithm was used with a minimum number of points as 15 and an epsilon radius of 10 nm, corresponding to the localization error, within Picasso render (*50*). The minimum number of points for DBSCAN was chosen as a parameter in accordance with the binding frequency of the imager strand and acquisition frame number. A custom-written MATLAB code was subsequently used for qPAINT processing (https://github.com/Simoncelli-lab/qPAINT_pipeline). Briefly, the sequence of dark times per cluster is compiled from localization time stamps and pooled for each cluster. A normalized cumulative histogram of dark times was then fitted with the exponential function 1 − exp(t/τd) to estimate the mean dark time, (τd), per cluster; which was then used to calculate the qPAINT index (qi). The protein number in each cluster was estimated by dividing qi/qi1, where qi1 was 0.5 x 10^2^ for both PSTPIP-1 and TCRζ images. Protein locations were mapped back to single molecule localizations using K-means clustering. An example representation of this qPAINT pipeline can be found in Fig. S6.

For protein clustering analysis, first, nearest neighbor distance (NND) analysis was used to compare protein distributions to complete spatial randomness (CSR), Fig. S8. The epsilon value for DBSCAN clustering was determined from the intersection of NND and CSR histograms (39 nm for PSTPIP-1 and 54 nm for TCRζ). This was compared to Ripley’s L function plots showing the L(r)-r over r function with positive deviations from expected CSR values (0) peaking at similar values. The minimum number of points for DBSCAN clustering of protein maps was given as 2, such that a cluster of proteins is defined as having at least two proteins. Protein clusters were quantified by the percentage of proteins in dimers, trimers, and larger clusters as presented in Fig. S9.

For two-color data, the association between PSTPIP-1 and TCRζ was analyzed by NND analysis of the first nearest neighbor from one protein to the other, compared to CSR maps at the same protein density. For this analysis, two CSR distributions were considered: one where the PSTPIP- 1 protein distribution was kept constant while TCRζ was randomized, and the other where the TCRζ distribution was fixed while PSTPIP-1 was randomized. Where significant associations closer than CSR were observed, then percentages and sizes of mixed clusters were calculated.

### Quantification and statistical analysis

Experiments used at least three independent cell passages, with a minimum of 10 cells per condition and two technical repeats. Statistical analyses and graph plotting was carried out in R (*49*). Shapiro-Wilk normality tests, histograms, and Q-Q plots were used to assess normality, guiding the choice of parametric or non-parametric tests. For two-sample comparisons, T-tests or Wilcoxon rank sum tests were applied. For multiple comparisons, ANOVA with Tukey’s HSD or Wilcoxon signed-rank tests with Bonferroni correction were used. Error bars represent SD (for means) or IQR (for boxplots). Statistical significance was defined as *p < 0.05, **p < 0.01, ***p < 0.001, ****p < 0.0001.

## Supplementary Materials

Fig. S1. F-actin in unstimulated WT and PTPN22 KO Jurkat T cells interacting with blank glass slides.

Fig S2. Actin remodeling in DMSO treated WT and PTPN22 KO Jurkat T cells interacting with aCD3/aCD28 coated slides.

Fig S3. PSTPIP-1 cytoskeletal localization in WT and PTPN22 KO Jurkat T cells inter- acting with aCD3/aCD28 coated slides.

Fig S4. Schematic representation of DNA-PAINT imaging and qPAINT principle targeting the TCRζ chain and PSTPIP-1 protein.

Fig S5. Localization precision of single molecule localizations for TCR and PSTPIP-1 DNA-PAINT imaging.

Fig S6. Schematic representation of the qPAINT analysis pipeline.

Fig S7. DNA-PAINT imaging and qPAINT quantification of TCR and PSTPIP-1 proteins coupled with SoRa super-resolution imaging of F-actin networks in unstimulated WT and PTPN22 KO Jurkat T cells.

Fig S8. 1st nearest neighbor distances (NND) of TCR and PSTPIP-1 proteins in aCD3/aCD28 activated and unstimulated WT and PTPN22 KO Jurkat T cells imaged with DNA-PAINT.

Fig S9. Percentage of TCR (A-B) and PSTPIP-1 (C-D) proteins in each protein cluster size in aCD3/aCD28 activated (A and C) and unstimulated (B and D) WT and PTPN22 KO Jurkat T cells imaged by DNA-PAINT.

Fig S10. TCR and PSTPIP-1 protein densities across the whole cell in aCD3/aCD28 activated and unstimulated WT and PTPN22 KO Jurkat T cells imaged by DNA-PAINT.

Videos S1-S2. Time-lapse imaging of actin remodeling in wild-type (WT) and PTPN22-knock- out (KO) Jurkat T cells interacting with aCD3/aCD28 coated slides. Related to Figure 1.

Videos S3-S4. Time-lapse imaging of actin remodeling in WT and PTPN22 KO Jurkat T cells pre-treated with the CK666 Arp 2/3 inhibitor interacting with aCD3/aCD28 coated slides. Related to Figure 2.

Video S5. Time-lapse imaging of actin remodeling in WT Jurkat T cells pre-treated with the LTV- 1 PTPN22 phosphatase action inhibitor interacting with aCD3/aCD28 coated slides. Related to Figure 4.

Videos S6-S9. Time-lapse imaging of actin remodeling in WT and PTPN22 KO Jurkat T cells interacting with high-affinity pTax pMHC or low-affinity pHuD pMHC coated slides.

## Acknowledgements

We sincerely thank Rose Zamoyska from Institute for Immunology and Infection Research, University of Edinburgh, Edinburgh, United Kingdom, for providing the WT and PTPN22 KO Jurkat T cell lines. We thank Andrew Vaughan and Ki Hng of the LMCB light microscopy facility for their assistance with the Nikon CSU-W1 SoRA. We also thank Robert Tetley from Nikon for his assistance with NIS-Elements software for the deconvolution of the images.

## Author contributions

Designed research: MDJ, MLD, APC, SS. Performed research: MDJ, CZ, OPLD. Contributed new reagents/analytic tools: MDJ, CZ, OPLD. Data analysis: MDJ, SS. Wrote the paper: MDJ, CZ, OPLD, MLD, APC, SS.

## Data and materials availability

All data needed to evaluate the conclusions in the paper are present in the paper or the Supplementary Materials. All original code has been deposited at https://github.com/Simoncelli-lab. Any additional information required to reanalyze the data reported in this paper is available from the corresponding author upon request.

