## Supplementary Materials for "PTPN22 Regulates T Cell Synapse Formation through PSTPIP1- Dependent Actin Remodeling"

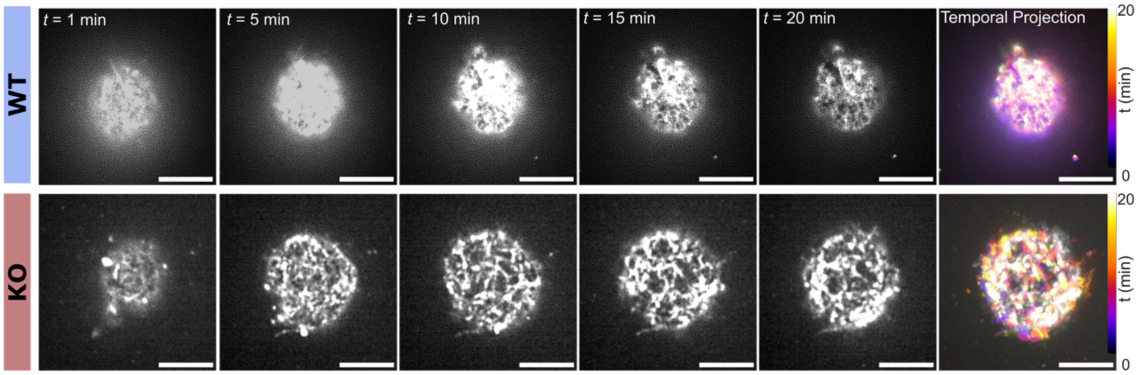


**Fig. S1. F-actin in unstimulated WT and PTPN22 KO Jurkat T cells interacting with blank glass slides.** Spinning disk confocal SoRa fluorescent imaging time-lapse of Jurkat T cells incubated with SiR actin from 1-20 minutes post settling. 1, 5-, 10-, 15-, and 20-minute time frames are shown. Temporal projections of spreading are depicted with pseudo-color images with color gradients transitioning from purple to white representing early to late time-points. Scale bar = 5 μm.

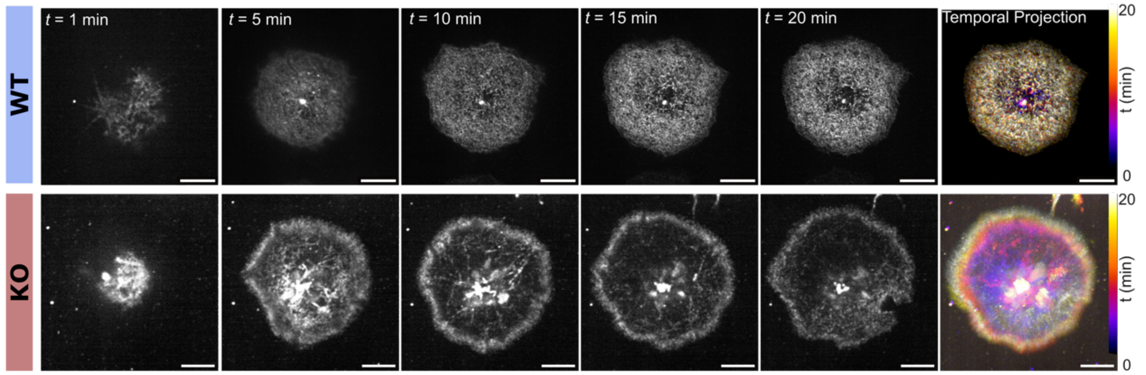


**Fig. S2. Actin remodeling in DMSO treated WT and PTPN22 KO Jurkat T cells interacting with aCD3/aCD28 coated slides.** Spinning disk confocal SoRa fluorescent imaging time-lapse of Jurkat T cells treated with DMSO and incubated with SiR actin from 1-20 minutes post settling. 1, 5-, 10-, 15-, and 20-minute time frames are shown. Temporal projections of spreading are depicted with pseudo-color images with color gradients transitioning from purple to white representing early to late time-points. Scale bar = 5 μm.

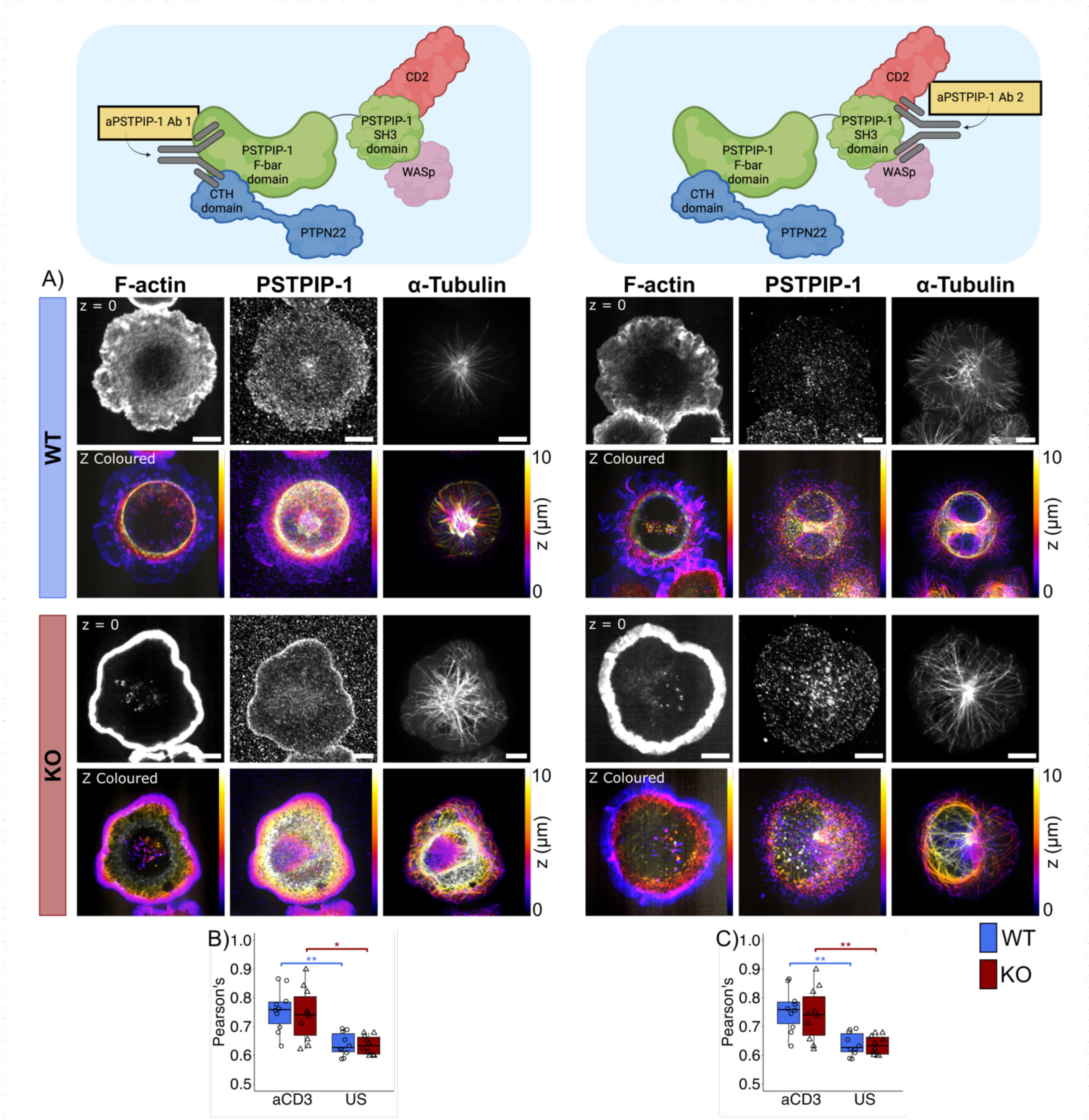


**Fig. S3. PSTPIP-1 cytoskeletal localization in WT and PTPN22 KO Jurkat T cells inter- acting with aCD3/aCD28 coated slides. (A)** 3D spinning disk confocal SoRa imaging of Jurkat T cells immunostained for the PSTPIP-1 protein, α-tubulin, and F-actin. Images were taken at 100 nm axial steps for 10 µm. PSTPIP-1 antibodies used are shown above each image respectively. The glass slide plane (z = 0) is shown, and axial projections are depicted with pseudo-color images with color gradients transitioning from purple to white representing low to high Z planes. Scale bar = 5 µm. **(B)** Pearson's correlation coefficient for the colocalization of PSTPIP-1 (Ab1) with actin and tubulin cytoskeletal structures in 3D. **(C)** Pearson's correlation coefficient for the colocalization of PSTPIP-1 (Ab1) with actin and tubulin cytoskeletal structures in 3D. For boxplots, n ≥ 10 cells from 3 independent passages. Statistical tests consisted of ANOVA with pairwise Tukey's HSD. Error bars on graphs represent interquartile range for boxplots and P values below 0.05 were considered significant using the following notation: * $p < 0.05$, ** $p < 0.01$, *** $p < 0.001$, **** $p < 0.0001$.

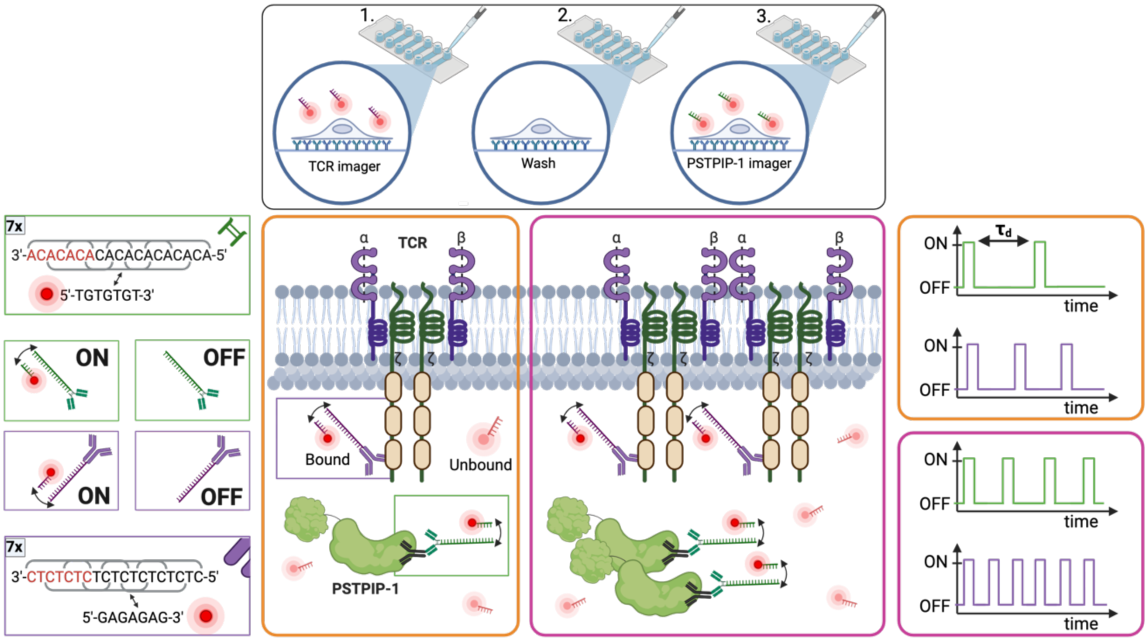


**Fig. S4. Schematic representation of DNA-PAINT imaging and qPAINT principle targeting the TCRζ chain and PSTPIP-1 protein.** Exchange PAINT methodology is depicted at the top with the first imager solution containing the imager strand complementary to the TCR docking strand added to image the TCR. This is then washed off in 2. The second imager solution containing the imager strand complementary to the PSTPIP-1 docking strand is then added to image PSTPIP-1. DNA sequences used are shown on the left-hand side with green corresponding to docking and imager strands used for PSTPIP-1 imaging and purple corresponding to docking and imager strands used for TCR imaging. 7x motifs show the repetitive nature of DNA strands used. Binding kinetics which control DNA-PAINT blinking kinetics are also shown here. Example TCR and PSTPIP-1 proteins are highlighted in the center with bound and unbound imager shown in the solution. The qPAINT principle is shown in the orange and pink boxes with one docking strand and corresponding ON-OFF traces in orange and two docking strands with corresponding ON-OFF traces shown in pink. This highlights the incrementally shorter dark time (τd) seen with increasing protein number. Created with BioRender.com

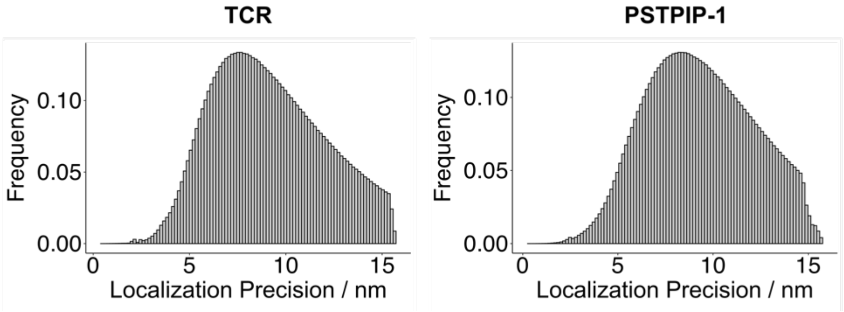


**Fig. S5. Localization precision of single molecule localizations for TCR and PSTPIP-1 DNA-PAINT imaging.**

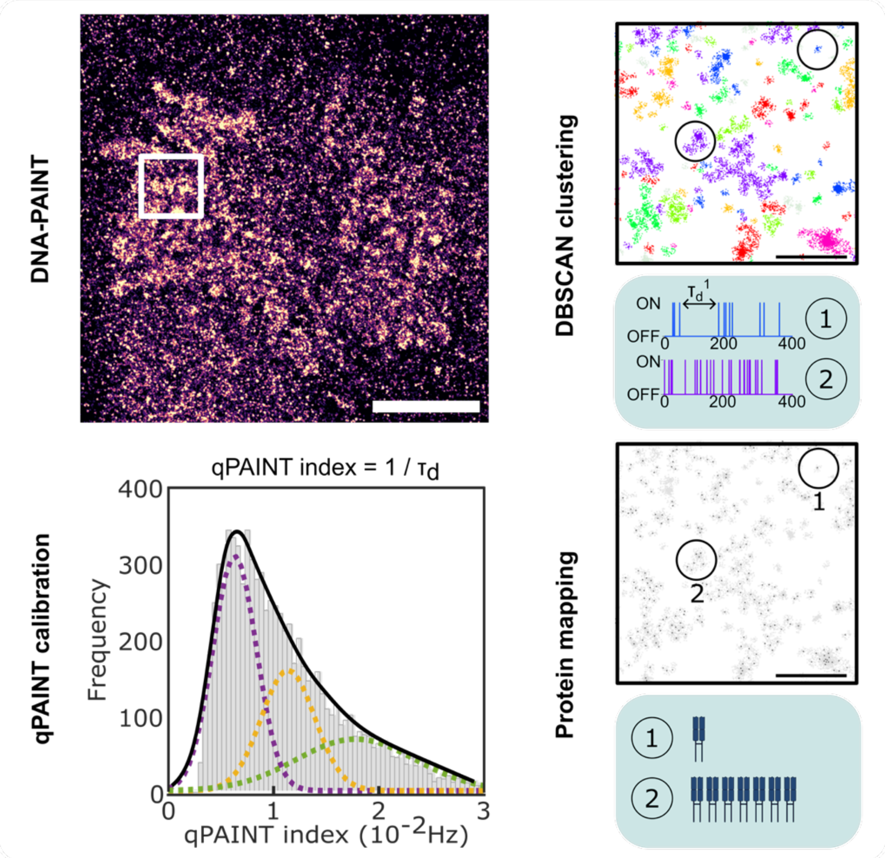


**Fig S6. Schematic representation of the qPAINT analysis pipeline.** Consisting of DNA-PAINT imaging, DBSCAN clustering of single molecule localizations, qPAINT calibration, protein quantification and mapping. DNA-PAINT scale bar = 1 µm, DBSCAN and protein map scale bar = 500 nm. Figure contains cartoon representations created with BioRender.com

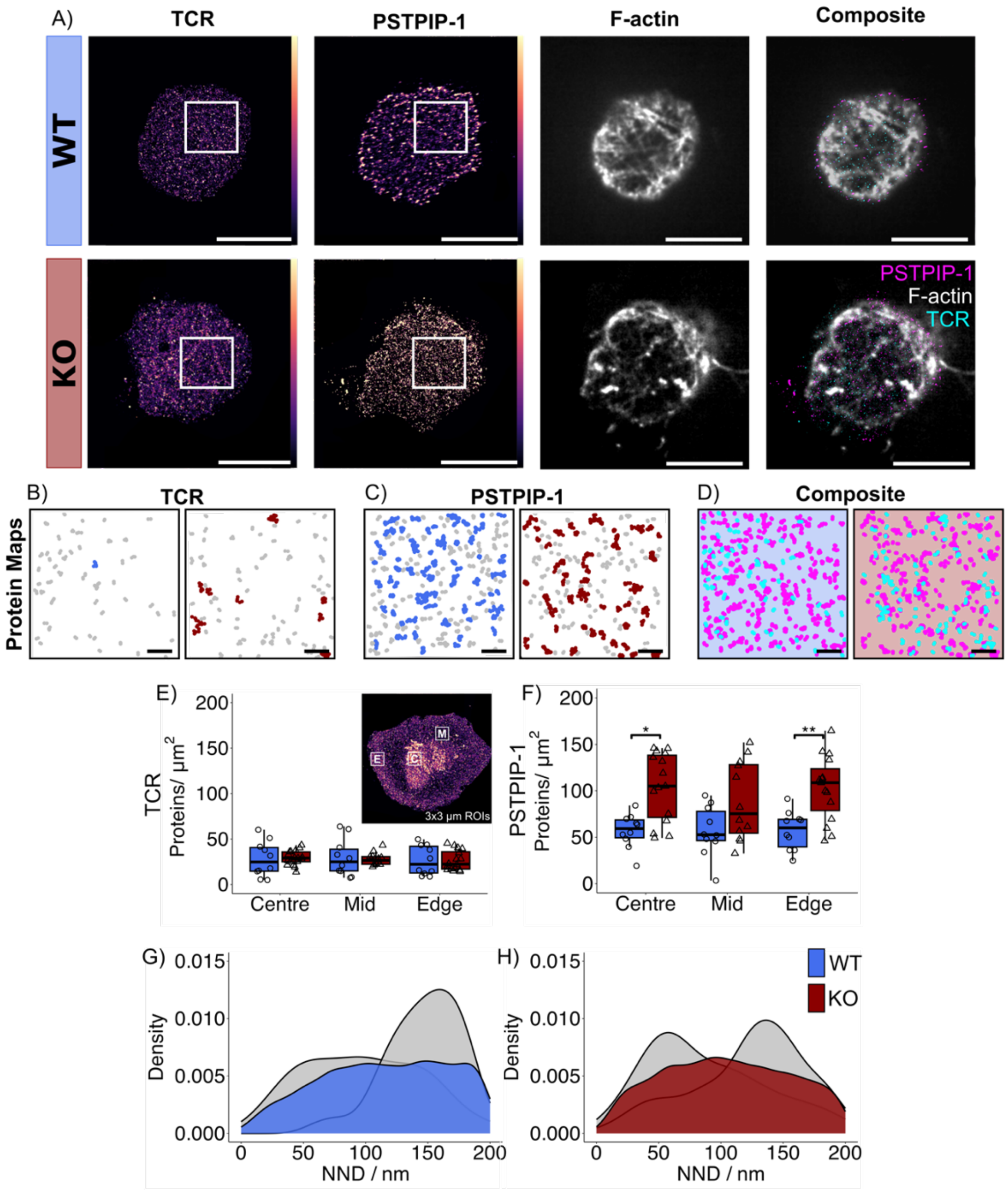


**Fig. S7. DNA-PAINT imaging and qPAINT quantification of TCR and PSTPIP-1 proteins coupled with SoRa super-resolution imaging of F-actin networks in unstimulated WT and PTPN22 KO Jurkat T cells. (A)** SoRa DNA-PAINT images of TCR and PSTPIP-1 coupled with fluorescent super-resolution images of F-actin in WT and PTPN22 KO Jurkat T cells after 15 minutes of interaction with PLL coated slides. DNA-PAINT images colored by density with yellow to purple representing high to low density of single molecule localizations. Scale bar = 5 µm. **(B)** Zoom region of TCRz protein maps calculated from the qPAINT pipeline as depicted by white boxes in **A**. Colored protein clusters contain ≥ 5 proteins. Scale bar = 500 nm. **(C)** Zoom region of PSTPIP-1 protein maps calculated from the qPAINT pipeline as depicted by white boxes in **A**. Colored protein clusters contain ≥ 5 proteins. Scale bar = 500 nm. **(D)** Composite overlay of TCR (cyan) and PSTPIP-1(magenta) protein maps. Scale bar = 500 nm. **(E)** TCR protein density in 3 x 3 µm$^2$ ROIs from the center, mid, and edge regions of unstimulated cells. **(F)** PSTPIP-1 protein density in 3 x 3 µm$^2$ ROIs from the center, mid, and edge regions of unstimulated cells. **(G)** Dual-color 1$^{st}$ nearest-neighbor distance (NND) analysis of PSTPIP-1 – TCR proteins in central ROIs of unstimulated WT cells compared to complete spatial randomness models (CSR) shown in grey.

**(H)** Dual-color 1st NND analysis of PSTPIP-1 – TCR proteins in central ROIs of unstimulated PTPN22 KO cells compared to CSR shown in grey. For all plots, $n \geq 10$ ROIs from $\geq 10$ cells from 3 independent passages. Statistical tests consisted of parametric ANOVA with pairwise Tukey's HSD for normally distributed data and non-parametric Wilcoxon signed-rank tests with Bonferroni correction for non-normally distributed data. Error bars represent interquartile range for boxplots and P values below 0.05 were considered significant using the following notation: * $p < 0.05$, ** $p < 0.01$, *** $p < 0.001$, **** $p < 0.0001$.

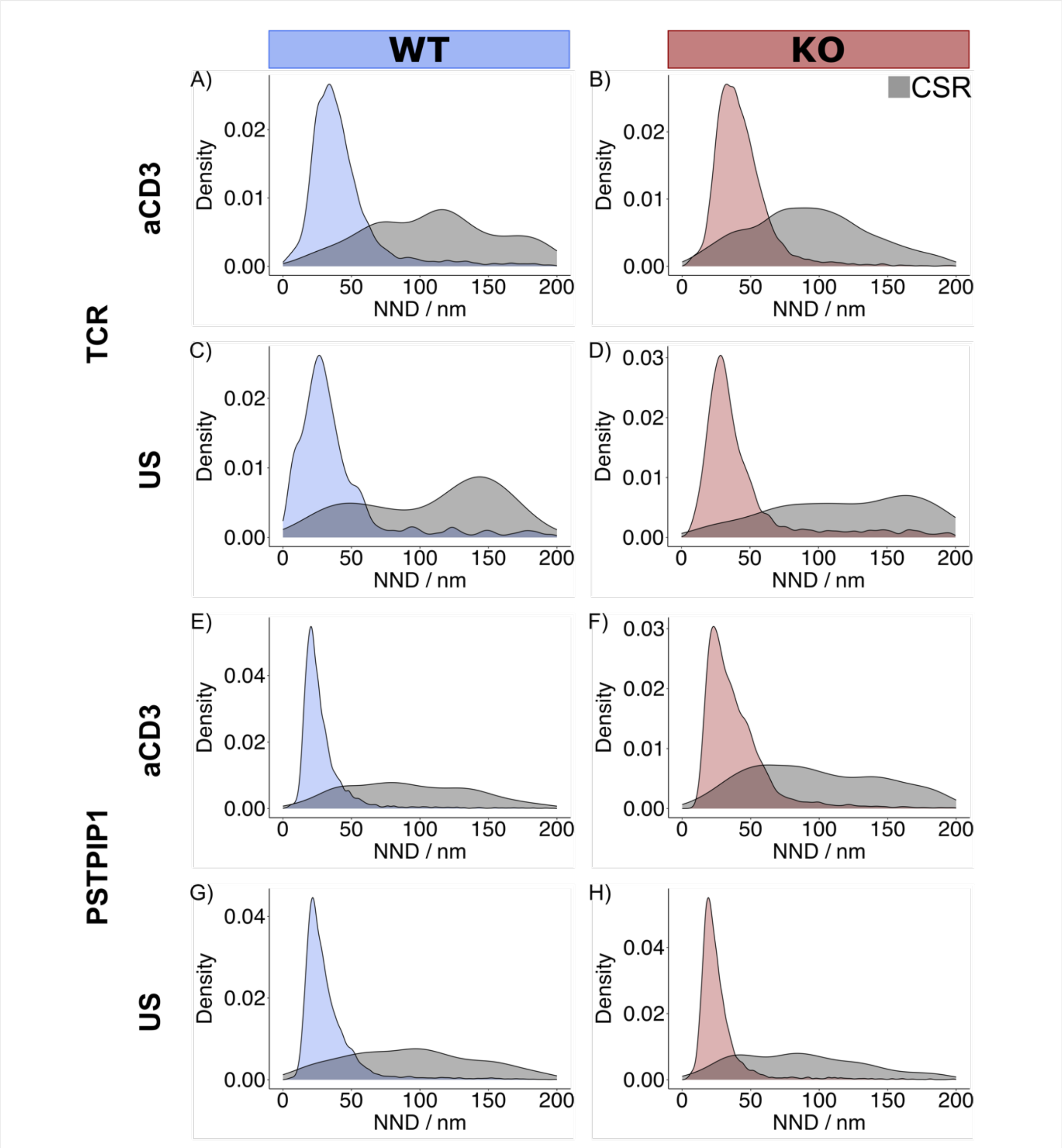


**Fig. S8. 1st nearest neighbor distances (NND) of TCR and PSTPIP-1 proteins in aCD3/aCD28 activated and unstimulated WT and PTPN22 KO Jurkat T cells imaged with DNA-PAINT.** Compared to complete spatial randomness (CSR). Distribution shown as probability density function. Data was pooled from all central ROIs in ≥ 10 cells from 3 independent passages.

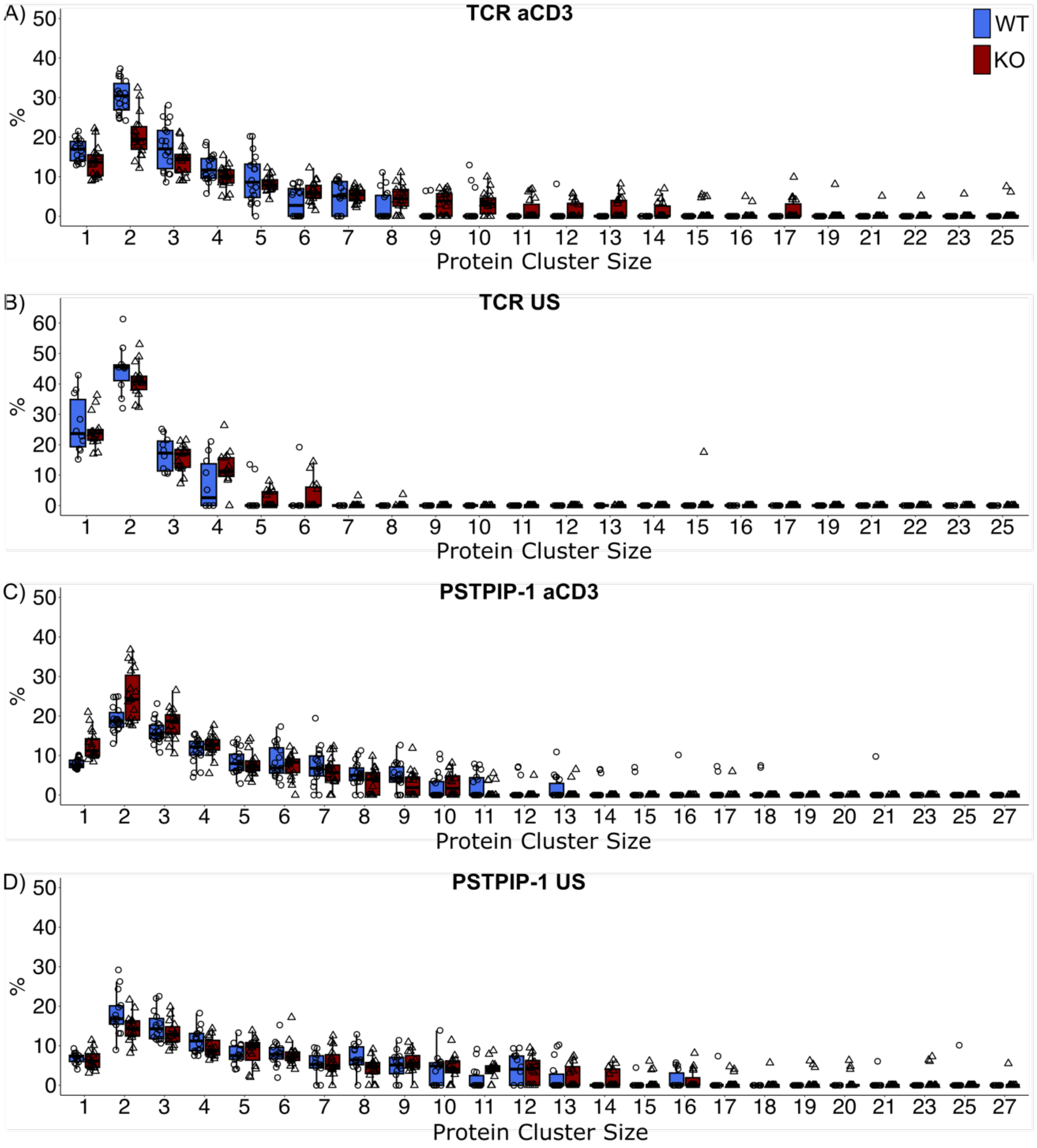


**Fig. S9. Percentage of TCR (A-B) and PSTPIP-1 (C-D) proteins in each protein cluster size in aCD3/aCD28 activated (A and C) and unstimulated (B and D) WT and PTPN22 KO Jurkat T cells imaged by DNA-PAINT.** Unstimulated represented as US. n = ROIs from ≥ 10 cells from 3 independent passages.

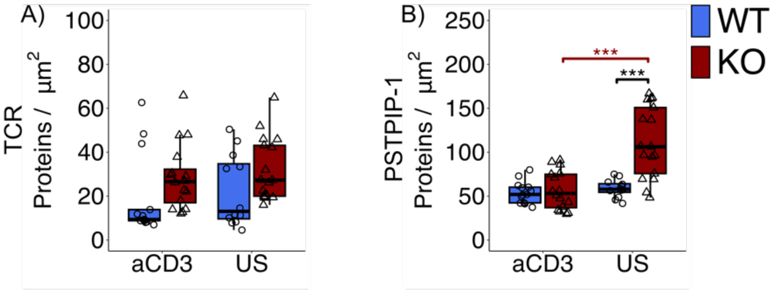


**Fig. S10. TCR and PSTPIP-1 protein densities across the whole cell in aCD3/aCD28 activated and unstimulated WT and PTPN22 KO Jurkat T cells imaged by DNA-PAINT.** Unstimulated represented as US. n ≥ 10 cells from 3 independent passages. Statistical tests consisted of parametric ANOVA with pairwise Tukey's HSD for normally distributed data and non-parametric Wilcoxon signed-rank tests with Bonferroni correction for non-normally distributed data. Error bars on graphs represent interquartile range, and P values below 0.05 were considered significant using the following notation: * $p < 0.05$, ** $p < 0.01$, *** $p < 0.001$, **** $p < 0.0001$.

**Video S1.** Time-lapse imaging of actin remodeling in wild-type (WT) Jurkat T cells interacting with aCD3/aCD28 coated slides. Images were captured for up to 20 minutes post-settling. Scale bar = 5 μm.

**Video S2**. Time-lapse imaging of actin remodeling in PTPN22 knock-out (KO) Jurkat T cells interacting with aCD3/aCD28 coated slides. Images were captured for up to 20 minutes post-settling. Scale bar = 5 μm.

**Video S3.** Time-lapse imaging of actin remodeling in WT Jurkat T cells pre-treated with the CK666 Arp 2/3 inhibitor interacting with aCD3/aCD28 coated slides. Images were captured for up to 20 minutes post-settling. Scale bar = 5 μm.

**Video S4**. Time-lapse imaging of actin remodeling in PTPN22 KO Jurkat T cells pre-treated with the CK666 Arp 2/3 inhibitor interacting with aCD3/aCD28 coated slides. Images were captured for up to 20 minutes post-settling. Scale bar = 5 μm.

**Video S5.** Time-lapse imaging of actin remodeling in WT Jurkat T cells pre-treated with the LTV-1 PTPN22 phosphatase action inhibitor interacting with aCD3/aCD28 coated slides. Images were captured for up to 20 minutes post-settling. Scale bar = 5 μm.

**Video S6.** Time-lapse imaging of actin remodeling in WT Jurkat T cells interacting with high-affinity pTax pMHC coated slides. Images were captured for up to 20 minutes post-settling. Scale bar = 5 μm.

**Video S7.** Time-lapse imaging of actin remodeling in PTPN22 KO Jurkat T cells interacting with high-affinity pTax pMHC coated slides. Images were captured for up to 20 minutes post-settling. Scale bar = 5 μm.

**Video S8.** Time-lapse imaging of actin remodeling in WT Jurkat T cells interacting with low-affinity pHuD pMHC coated slides. Images were captured for up to 20 minutes post-settling. Scale bar = 5 μm.

**Video S9.** Time-lapse imaging of actin remodeling in PTPN22 KO Jurkat T cells interacting with low-affinity pHuD pMHC coated slides. Images were captured for up to 20 minutes post-settling. Scale bar = 5 μm.
